# Mitotic chromatin compaction tethers extrachromosomal DNA to chromosomes and prevents their mis-segregation into micronuclei

**DOI:** 10.1101/2025.08.21.671584

**Authors:** Lu M. Yang

## Abstract

Extrachromosomal DNA (ecDNA) is linked to aggressive cancer growth, treatment resistance, and shorter survival across a wide variety of cancers. ecDNA promotes intratumoral genetic heterogeneity, enhanced oncogene expression, and accelerated tumor evolution, driving tumor pathogenesis. ecDNA lack centromeres and segregate to daughter cell nuclei during mitosis by tethering to chromosomes. However, the mechanisms involved in this tethering are incompletely understood. Here, I present evidence that ecDNA tethering to chromosomes is coupled to chromatin compaction during mitotic chromosome formation, which acts to generally increase chromatin-chromatin interaction. Using a cancer cell line model, I show that decompacting mitotic chromatin under hypotonic conditions and by increasing histone acetylation untethers ecDNA from chromosomes, leading to their mis-segregation into micronuclei after mitosis. Additionally, overexpression of the mitotic chromosome surfactant Ki67 untethers ecDNA from chromosomes, leading to their mis-segregation into micronuclei. These findings show that the mechanisms involved in chromatin compaction are important for tethering ecDNA to chromosomes and preventing their mis-segregation into micronuclei. I propose a model in which interactions between ecDNA chromatin fibers and chromosomal chromatin contribute to ecDNA segregation into daughter cells during cell division.

## Introduction

Focal amplifications involving proto-oncogenes are a hallmark of cancer(Albertson, 2006; Schwab, 1998; Stratton et al., 2009). Focal amplifications can be located either on chromosomes, often forming homogeneously staining regions (HSRs), or on extrachromosomal DNA (ecDNA), traditionally known as double minutes (DMs)(Storlazzi et al., 2010). ecDNA are chromatinized, large (ranging from 100s of kb to several Mb in size), circular, and acentric(Carroll et al., 1987; Haaf & Schmid, 1988; Levan & Levan, 1978; Levan et al., 1976; Turner et al., 2017; Von Hoff et al., 1988; Wahl, 1989). Compared to HSRs and other forms of focal amplifications, ecDNA-containing cancers are associated with lower patient survival(Kim et al., 2020; Pongor et al., 2023). Importantly, ecDNA lack centromeres and tether to either sister chromatid during mitosis to ensure that they are segregated into daughter cell nuclei(Jack et al., 1987; Kanda et al., 2001; Levan & Levan, 1978) (Fig 1a). This stochastic manner of inheritance leads to unequal ecDNA segregation, thereby contributing to cancer heterogeneity(Chapman et al., 2023; deCarvalho et al., 2018; Fiorini et al., 2025; Kanda et al., 1998; Lange et al., 2022; Lundberg et al., 2008; Turner et al., 2017). While first described in 1978, the mechanism underlying how ecDNA tether to mitotic chromosomes is only beginning to be elucidated(Ilić et al., 2022). Recent studies suggest that mitotic ecDNA transcription(Nichols et al., 2025; Xie et al., 2025) has a role in mediating ecDNA tethering. I hypothesized that the molecular mechanisms that mediate the organization and compaction of mitotic chromosomes, acting as a general chromatin-attractive force to increase chromatin-chromatin interaction, also mediate ecDNA tethering.

**Figure 1.**
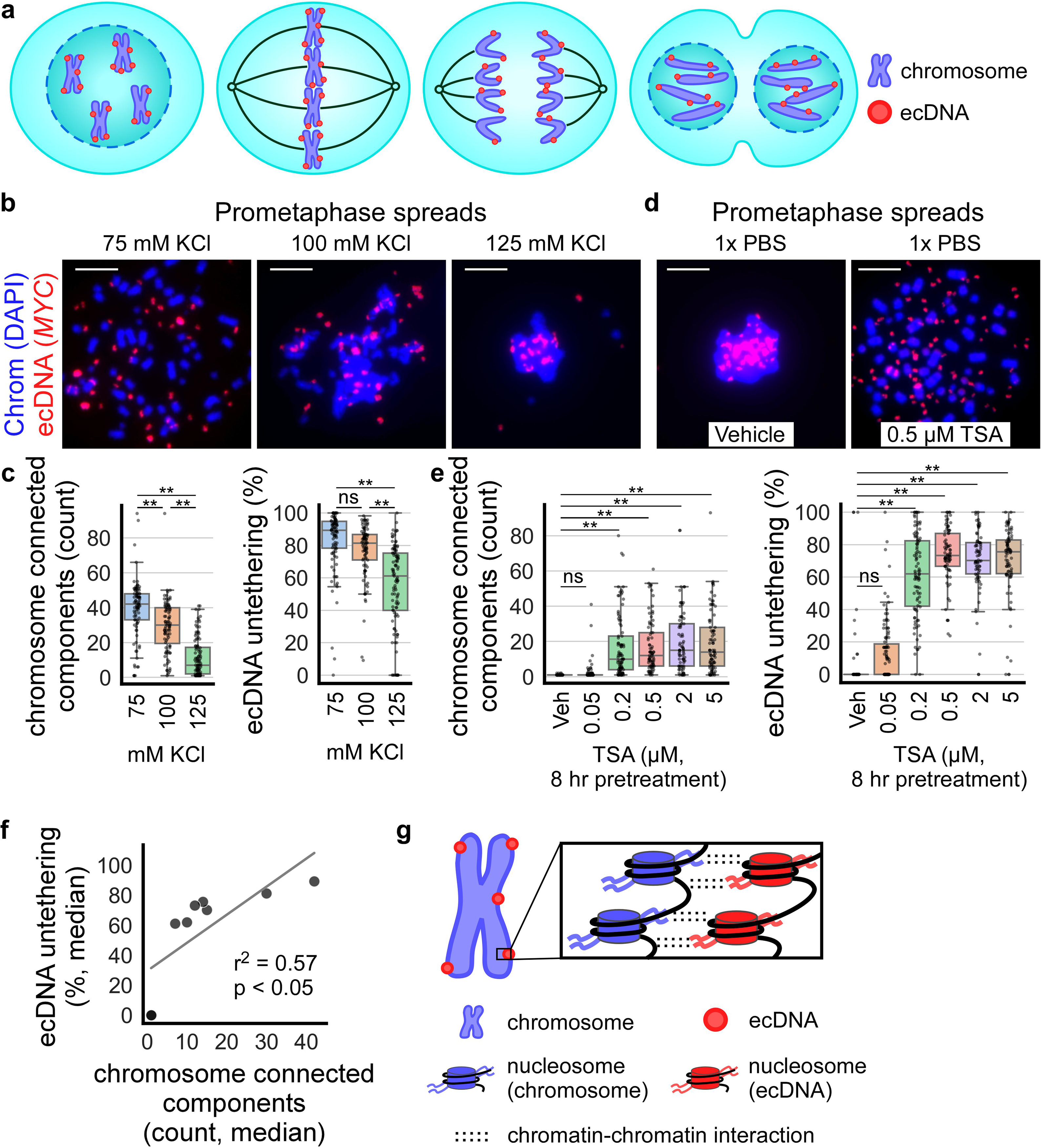
The prometaphase spread technique untethers ecDNA and chromosomes by decompacting mitotic chromatin. **a)** Schematic of an ecDNA-containing cell undergoing mitosis (from left to right: prophase, metaphase, anaphase, telophase). Acentric ecDNA (red) have been observed to tether to or “hitchhike” on chromosomes (blue) during mitosis to ensure their proper segregation and inheritance by daughter cells. **b)** Prometaphase spreads performed on colcemid-arrested COLO320DM cells with the conventional hypotonic solution (75 mM KCl) and higher osmolarity solutions (100 mM and 125 mM KCl), producing varying amounts of chromosome (DAPI, blue) individualization and ecDNA (*MYC* FISH, red) untethering. **c)** Quantification of panel b. Left: boxplots quantifying chromosome individualization in prometaphase spreads performed with varying solution osmolarity; from left to right, n=3, 3, 3 biological replicates and 95, 92, 105 cells; one-way ANOVA, F=100.7, p<.001; **p<.01 by Tukey’s honest significant difference (HSD), ns = not significant. Right: quantification of ecDNA untethering; one-way ANOVA, F=48.3, p<.001. Chromosome individualization is represented by the number of connected components identified as chromosomes by ecSeg. ecDNA untethering is represented by the number of ecDNA unattached to chromosomes divided by the total number of ecDNA not completely surrounded by chromosomes. **d)** Prometaphase spreads performed with incubation in 1x PBS using COLO320DM cells pretreated for 8 hr with vehicle (0.1% DMSO) or varying concentrations of Trichostatin A (TSA). **e)** Quantification of panel d. Left: quantification of chromosome individualization in prometaphase spreads performed with 1x PBS on cells pretreated with TSA; n=3, 3, 3, 3, 3, 3 biological replicates and 88, 113, 96, 80, 76, 84 cells; one-way ANOVA, F=38.5, p<.001; **p<.01 by Tukey’s HSD, ns = not significant. Right: quantification of ecDNA untethering; one-way ANOVA, F=166.5, p<.001. **f)** Linear regression analysis of median chromosome individualization vs. ecDNA untethering of prometaphase spread conditions in panels b-e (n=9). **g)** Schematic: electrostatic/hydrophobic mitotic compaction forces at the level of nucleosomes tether ecDNA (red) to mitotic chromosomes (blue). In all panels, scale bar = 10 µm.

During cell division, mitotic spindles associate with chromosomes at centromeres to direct chromosomal segregation into daughter cell nuclei. Chromosomes form during mitosis and move along the mitotic spindles, ensuring the segregation of one genome copy into each of the daughter cell(Spicer & Gerlich, 2023). They are discrete, cylindrical structures composed of densely packed chromatin, organized through a series of coordinated molecular mechanisms, including 1) topological organization of chromatin into loops with helical periodicity; 2) global chromatin compaction; and 3) the regulation of chromosome surface by surfactants such as Ki67(Spicer & Gerlich, 2023). *Mechanism 1:* the typical chromosome cylindrical form arises from chromatin loops organized around the chromosome axis by molecular motors such as condensin that mediate DNA loop extrusion(Davidson & Peters, 2021; Ganji et al., 2018; Gibcus et al., 2018; Hirano, 2016; Paulson et al., 2021; Samejima et al., 2025). These molecular motors help stiffen chromosomes(Gerlich et al., 2006; Houlard et al., 2015; Ribeiro et al., 2009; Sun et al., 2018) and shape their mitotic structure(Beckwith et al., 2025; Cuylen et al., 2011; Hirano & Mitchison, 1994; Hirota et al., 2004; Ono et al., 2003). *Mechanism 2:* global chromatin compaction is driven by a network of dynamic and weak interactions between nucleosomes and chromatin-associated proteins that reduces chromatin volume(Spicer & Gerlich, 2023). This process consists in the folding and local compaction of the 10 nm fiber of nucleosomes, giving rise to denser and irregular chromatin structure. Such global chromatin compaction is distinct from topological organization of mitotic chromosomes by condensin and other molecular motors(Beckwith et al., 2025; Hibino et al., 2024; Spicer & Gerlich, 2023; Strickfaden et al., 2020) and is regulated by histone deacetylation and free intracellular Mg^2+^ concentration that modulate chromatin-chromatin affinity interactions during mitosis(Cimini et al., 2003; Kruhlak et al., 2001; Maeshima et al., 2018; Schneider et al., 2022; Wilkins et al., 2014). *Mechanism 3:* factors that regulate the chromosome surface mediate the spatial separation of chromatin into distinct chromosome bodies in late prophase, which prevents chromosome coalescence and enables spindle microtubules to access the kinetochores(Spicer & Gerlich, 2023). The protein Ki67 has been shown to act as chromosome surfactant by forming a repulsive layer of proteins and RNA at the periphery of mitotic chromosomes, thereby promoting chromosome individualization(Booth et al., 2014; Cuylen et al., 2016; Saiwaki et al., 2005; Takagi et al., 2018).

Here, I use an ecDNA-containing cancer cell line to study the role of mitotic chromatin compaction on ecDNA tethering to chromosomes and their mitotic inheritance. I focus on disrupting mechanisms 2 and 3 via small molecule drug treatment, cell osmolarity alteration, and genetic manipulation. I show that mitotic chromatin decompaction, induced either by hypotonic conditions or histone deacetylase (HDAC) inhibition, disrupts ecDNA-chromosome tethering. Additionally, I identify Ki67, which coats the surface of both mitotic chromosomes(Booth et al., 2014; Cuylen et al., 2016; Saiwaki et al., 2005; Takagi et al., 2018) and ecDNA, as a modulator of ecDNA tethering. Untethered ecDNA have an increased likelihood of mis-segregating into micronuclei. I propose that global chromatin compaction, driven by increased chromatin-chromatin interactions (mechanism 2), contributes to ecDNA tethering by promoting chromatin interactions between ecDNA and chromosome. Furthermore, I propose that the surfactant properties of Ki67 (mechanism 3) control ecDNA tethering by coating the surface of both ecDNA and chromosomes.

## Results

### The prometaphase spread technique untethers ecDNA and chromosomes by decompacting mitotic chromatin

COLO320DM cells are a colorectal cancer cell line containing *MYC* amplification on ecDNA, which can be visualized using *MYC* fluorescence in situ hybridization (FISH). On prometaphase spreads of these cells, ecDNA appear spatially dissociated from the chromosomes, suggesting that they are untethered (example: Fig 1b, leftmost panel). This untethering has also been observed in prometaphase spreads of other cell lines containing ecDNA(Turner et al., 2017). I reasoned that ecDNA untethering could be linked to the reduction of intracellular ion concentration caused by incubation in hypotonic solution during prometaphase spread preparations (extended Fig 1a), which may disrupt interactions involved in mitotic chromatin compaction(Brinkley et al., 1980; Gibson et al., 2019; Hansen et al., 2021; Iida et al., 2024).

To test this, I performed prometaphase spreads using higher osmolarity variations (100 mM and 125 mM KCl) of the conventional 75 mM KCl hypotonic solution (Fig 1b). 100 mM and 125 mM KCl are still hypotonic relative to the cells, but the net inflow of water into cells is expected to be decreased compared to 75 mM KCl. Both higher osmolarity treatments significantly reduced chromosome individualization, as measured by the number of chromosome connected components detected using ecSeg(Rajkumar et al., 2019), from 39.2 ± 16.8 (mean ± standard deviation) with 75 mM KCl to 29.3 ± 14.6 and 11.4 ± 11.0 with 100 mM and 125 mM KCl, respectively. Similarly, ecDNA untethering decreased with the higher osmolarity treatments, from 83.3% ± 18.6 with 75 mM KCl to 76.7% ± 16.9 and 55.4% ± 25.8 with 100 mM and 125 mM KCl, respectively (Fig 1b,c). These observations suggest that ionic strength modulates ecDNA tethering to chromosomes, possibly via decompacting mitotic chromatin.

In addition to ionic strength, chromatin compaction is modulated by histone tail acetylation(Gibson et al., 2019; Wilkins et al., 2014; Zhiteneva et al., 2017). To assess whether acetylation modulates chromosome individualization and ecDNA untethering similarly to ionic strength, I treated cells with Trichostatin A (TSA), a pan-histone deacetylase (HDAC) inhibitor that causes histone hyperacetylation and decompacts mitotic chromatin(Cimini et al., 2003; Görisch et al., 2005; Kruhlak et al., 2001; Schneider et al., 2022; Strickfaden et al., 2020). I treated COLO320DM cells for 8 hr with varying concentrations of TSA before performing prometaphase spreads in 1x PBS rather than 75 mM KCl (Fig 1d). Staining for pan-Histone H3 acetylation confirmed an increase in acetylation on both ecDNA and chromosomes in cells treated with TSA in a dose-dependent manner (extended Fig 1b). TSA-treated cells presented both increased chromosome individualization (17.3 ± 14.9 in 0.5 µM TSA-treated cells) and ecDNA untethering (73.5% ± 17.3 in 0.5 µM TSA-treated cells) compared to vehicle-treated cells (1.0 ± 0.2 chromosome individualization and 7.2% ± 22.8 ecDNA untethering) (Fig 1d,e). This indicates that histone hyperacetylation disrupts ecDNA tethering to chromosomes, possibly via decompacting mitotic chromatin.

Notably, under hypotonic conditions and in TSA-treated cells, ecDNA untethering correlates with chromosome individualization in prometaphase spreads (Fig 1f), suggesting that both chromosome-chromosome and ecDNA-chromosome tethering are regulated by ionic strength and histone acetylation (Fig 1g). These observations imply that chromosome individualization and ecDNA tethering may be governed by the interactions involved in chromatin compaction.

### Hypotonic conditions and HDAC inhibition untether ecDNA

While prometaphase spreads allow visualization of separated chromosomes, they can alter nuclear organization. To further assess the link between chromatin compaction and ecDNA tethering in a more physiologically relevant context, I evaluated ecDNA tethering in COLO320DM metaphase cells cultured and fixed on glass coverslips. In these preparations, chromosomes are aligned at the metaphase plate, allowing the visualization of untethered ecDNA that detach from the metaphase plate while better preserving spatial relationships within the cell. To test the effects of variation in ionic conditions on ecDNA tethering, I treated cells for 15 min with solutions at five different ionic strengths: i. 1x media (isotonic control), ii. 0.75x media (hypotonic), iii. 0.5x media (hypotonic), iv. 1:1 mix of 1x PBS with 1x media (1x PBS-media, relatively isotonic control), and v.

1:1 mix of 1.5x PBS with 1x media (1.25x PBS-media, hypertonic). After treatment, cells were fixed for 10 min in paraformaldehyde dissolved in 1x, 0.75x, 0.5x, or 1.25x PBS matching the osmolarity of the incubation media to preserve the ionic conditions of the treatment. Cells were stained with DAPI to visualize DNA and *MYC* FISH to visualize ecDNA. ecDNA untethering significantly increased in cells incubated in hypotonic 0.75x (22.7% ± 15.2 untethered, mean ± standard deviation) and 0.5x (38.9% ± 17.1) media compared to 1x media (5.5% ± 6.0) (Fig 2a,b). Conversely, ecDNA untethering slightly but statistically significantly decreased in cells incubated in hypertonic 1.25x PBS-media (6.3% ± 6.8) compared to 1x PBS-media (8.3% ± 9.2). The effects of ionic strength on ecDNA tethering are consistent with known effects of ionic strength on chromatin compaction: low salt conditions induce chromatin decompaction by reducing molecular interactions, while higher salt concentrations promote compaction(Brinkley et al., 1980; Gibson et al., 2019; Hansen et al., 2021; Iida et al., 2024).

**Figure 2.**
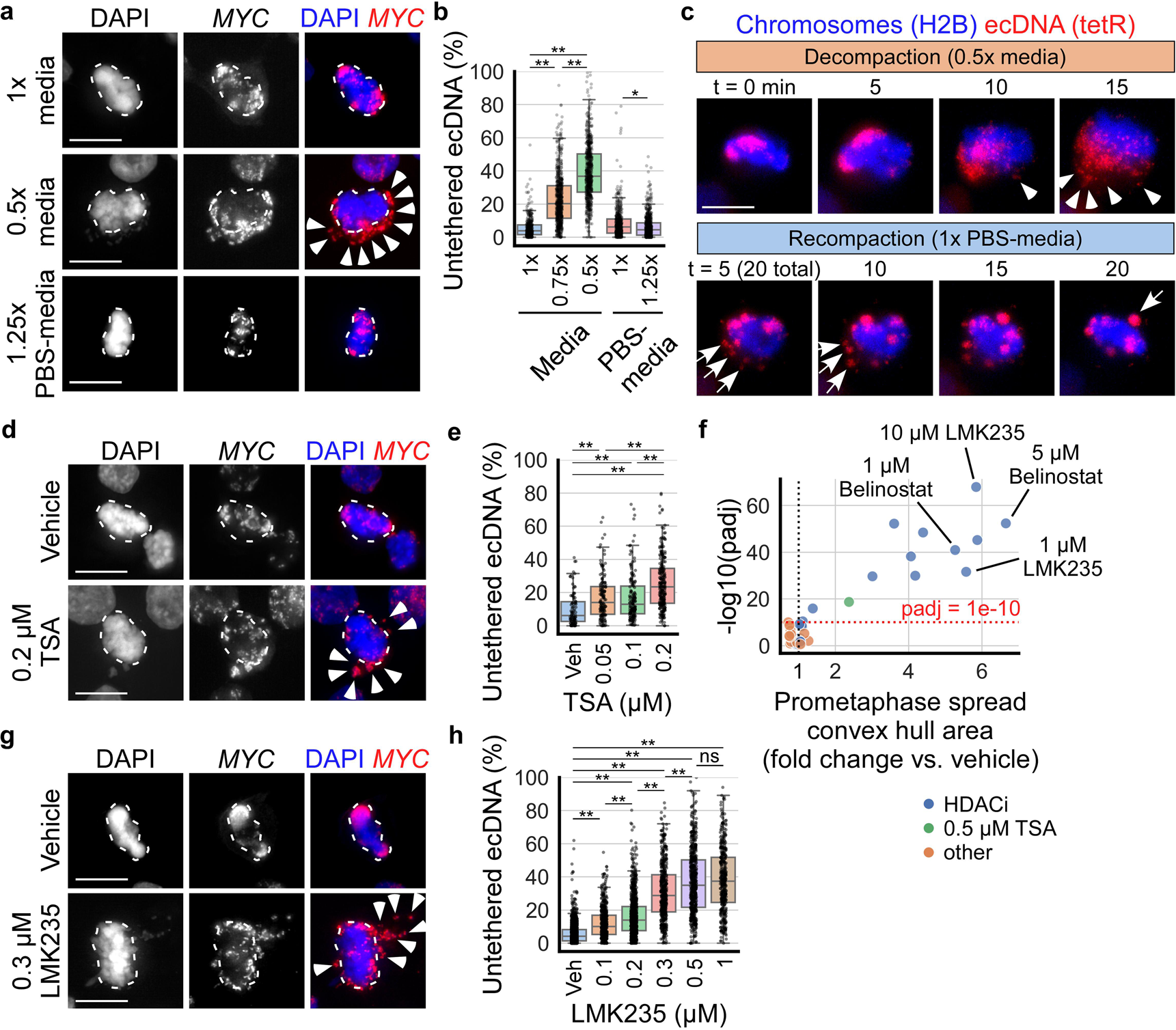
Hypotonic conditions and HDAC inhibition untether ecDNA. **a)** Fixed metaphase COLO320DM cells cultured on glass coverslips treated for 15 min with 1x media, 0.75x media (not shown), 0.5x media, 1:1 mix of 1x PBS with 1x media (1x PBS-media, not shown), and 1:1 mix of 1.5x PBS with 1x media (1.25x PBS-media); dashed outlines indicate metaphase plate chromosomes (DAPI, blue), arrowheads indicate untethered ecDNA (identified by *MYC* FISH signal, red). **b)** Boxplots quantifying untethered ecDNA per cell from panel a, represented by % of *MYC* FISH signal in a cell unattached to chromosomes aligned at the metaphase plate (non-overlapping pixels); from left to right, n=4, 6, 6, 7, 4 biological replicates and 427, 672, 691, 532, 686 cells; one-way ANOVA, F=882.3, p<.001; *p<.05, **p<.01 by Tukey’s HSD. **c)** Live imaging of COLO320DM cells expressing H2B-emiRFP670 (chromatin, blue) and tetR-mNeonGreen which binds to a tetO-96mer repeat sequence inserted near *MYC* loci (*MYC*, red). Cells were arrested at metaphase with 10 µM MG132(Wee et al., 2022) and placed in hypotonic 0.5x media at t=0 min to decompact chromatin; arrows indicate untethered ecDNA; after image acquisition at t=15 min, cells were placed in relatively normotonic 1x PBS-media to recompact chromatin; arrows indicate ecDNA-ecDNA tethering. Representative images of n=5 biological replicates and >50 cells. **d)** Fixed metaphase COLO320DM cells cultured on glass coverslips treated for 8 hr with vehicle (0.1% DMSO) or indicated concentrations of TSA; dashed outlines indicate metaphase plate chromosomes (DAPI, blue), arrowheads indicate untethered ecDNA (identified by *MYC* FISH signal, red). **e)** Boxplots quantifying untethered ecDNA per cell from panel d; from left to right, n=3, 3, 3, 3 biological replicates and 98, 144, 161, 246 cells; one-way ANOVA, F=34.9, p<.001; **p<.01 by Tukey’s HSD. **f)** Modified volcano plot summarizing the results of the targeted screen for drugs with the ability to untether ecDNA and chromosomes, quantified by the area of the convex hull of the prometaphase spread produced without hypotonic solution incubation. TSA (0.5 µM) was included as positive control. padj = adjusted Student’s t-test p value using Bonferroni multiple test correction (44 total comparisons were made). **g)** Metaphase COLO320DM cells cultured on glass coverslips treated for 24 hr with vehicle (0.1% DMSO) or indicated concentrations of LMK235; dashed outlines indicate metaphase plate chromosomes, arrowheads indicate untethered ecDNA. **h)** Quantification of untethered ecDNA per cell from panel g; n=11, 3, 7, 4, 3, 3 biological replicates and 1278, 501, 875, 472, 571, 388 cells; one-way ANOVA, F=735.1, p<.001; **p<.01 by Tukey’s HSD, ns = not significant. In all panels, scale bar = 10 µm.

Prior studies have suggested that DNA double-strand breaks (DSBs) may cause ecDNA untethering, aggregation, and mis-segregation(Oobatake & Shimizu, 2020; Shimizu et al., 2007; Tanaka & Shimizu, 2000). To determine whether hypotonic and hypertonic treatments alter DSB levels, I examined by IF the expression of histone H2A.X phosphorylation (pH2A.X), a marker of DSB(Rogakou et al., 1999) which peaks at 2-3 min after DSB formation(Giunta et al., 2010; Trakarnphornsombat & Kimura, 2023). I found that while 0.5x hypotonic media incubation slightly decreased the number of pH2A.X foci per metaphase cell (extended Fig 2a,b), in every condition, the number of pH2A.X foci did not correlate with the extent of ecDNA untethering (extended Fig 2c). This suggests that incubation in hypotonic media does not induce ecDNA untethering by promoting DSBs.

Next, I reasoned that mitotic chromatin in a hypercompacted state will be more resistant to ecDNA untethering by hypotonic incubation, if the mechanism of untethering is via chromatin decompaction. Treatment of mitotic cells with 10 mM sodium azide (NaN_3_) and 50 mM 2-Deoxy-D-glucose (2-DG) has been shown to hypercompact chromatin by increasing the intracellular free Mg^2+^ concentration available to associate with chromatin(Maeshima et al., 2018). I treated cells for 3 hr with 10 mM NaN_3_ + 50 mM 2-DG dissolved directly into the cell culture media, followed by a 15 min incubation in 1x, 0.75x, or 0.5x media. Chromatin hypercompaction by NaN_3_ + 2-DG significantly reduced the amount of ecDNA untethering by hypotonic 0.75x and 0.5x media (extended Fig 2d,e). Altogether, these results show that intracellular ionic strength regulates ecDNA tethering by modulating chromatin compaction.

To examine ecDNA untethering in real time, I employed live cell imaging using a COLO320DM cell line genetically modified to express histone H2B-emiRFP670 for chromatin visualization, and tetR-mNeonGreen that binds to a Tet-operator (tetO) array inserted onto ecDNA for ecDNA visualization(Hung et al., 2024). Cells were arrested at metaphase with MG132(Wee et al., 2022) and placed in hypotonic 0.5x media (Fig 2c). After 5-10 min of incubation in 0.5x media, ecDNA began moving away from the chromosomes aligned at the metaphase plate. At 15 min, most of the ecDNA appeared as smaller scattered foci, suggesting they have untethered from chromosomes and each other. To test for the reversibility of the untethering process, after 15 min in hypotonic 0.5x media, cells were placed back into isotonic conditions (1x PBS-media). I observed ecDNA compacting into larger foci and by 20 min in isotonic conditions most of the ecDNA appeared retethered to the chromosomes or with each other (Fig 2c). These results suggest that tethering and untethering are reversible processes, like chromatin compaction regulation by intracellular ionic strength(Hansen et al., 2021).

As described above, I found that treatment with the HDAC inhibitor TSA leads to untethering of ecDNA from chromosomes in prometaphase spreads. I confirmed this finding using COLO320DM metaphase cells cultured and fixed on glass coverslips, which, when treated with varying concentrations of TSA for 8 hr, showed increased ecDNA untethering compared to vehicle-treated cells (9.9% ± 10.0 untethered, mean ± standard deviation): 17.2% ±13.2 untethered with 0.05 µM TSA, 17.1% ±14.4 with 0.1 µM, and 25.9% ± 15.7 with 0.2 µM; above 0.2 µM, I was unable to find many cells that progressed to metaphase (Fig 2d,e).

TSA is a broad-spectrum HDAC inhibitor blocking class I and II HDACs. To determine whether other HDAC inhibitors may similarly regulate ecDNA tethering to chromosomes, I performed a targeted screen of general and class-specific HDAC inhibitors (HDACi). Twelve HDACi were tested, including Belinostat and Abexinostat (pan-HDACi(Ho et al., 2020)), Fimepinostat (HDAC class I and IIb inhibitor), Pracinostat (HDAC class I and II inhibitor), Mocetinostat and Entinostat (HDAC class I inhibitor), LMK235 and TMP269 (HDAC class IIa inhibitor), BRD73954 (HDAC class IIb inhibitor), Tasquinimod (HDAC4 inhibitor), RGFP966 (HDAC3 inhibitor(Gandhi et al., 2022)), and Santacruzamate A (HDAC2 inhibitor(Gandhi et al., 2022). Additionally, I included nine non-HDACs inhibitor drugs with targets suggested to be involved in chromatin compaction through different mechanisms, such as Nicotinamide (pan sirtuin inhibitor(Gandhi et al., 2022)), EX527 (SIRT1 inhibitor(Gandhi et al., 2022)), CTPB (p300/CBP histone acetyltransferase activator(Balasubramanyam et al., 2003; Gandhi et al., 2022)), RRx001 (DNA methyltransferase inhibitor(Zhao et al., 2015)), 5-Azacytidine (DNA methyltransferase inhibitor(Djeghloul et al., 2023)), GSK343 (EZH2 inhibitor(Djeghloul et al., 2023)), ZM447439 and Hesperadin (Aurora kinase inhibitor(Gadea & Ruderman, 2005; Hauf et al., 2003)), and Paprotrain (MKLP-2 inhibitor(Orr et al., 2021)).

I treated COLO320DM cells with each of these drugs for 24 hr at multiple concentrations ranges and performed prometaphase spreads using incubation in 1x PBS rather than the conventional hypotonic solution. I then quantified the area of the convex hull of prometaphase spreads (extended Fig 2f) as an approximation of ecDNA and chromosome untethering. I found that only inhibitors of HDAC classes I and II effectively untethered ecDNA and individualized chromosomes, including Belinostat, Abexinostat, Fimepinostat, Pracinostat, and LMK235 (Fig 2f, extended Fig 2g). These effects were confirmed by treating COLO320DM cells cultured on glass coverslips with LMK235 or Belinostat for 24 hr, which untethered ecDNA in a dose-dependent manner (Fig 2g,h and extended Fig 2h,i). I investigated the possibility of DSB formation as a confounding variable by examining pH2A.X IF and observed no correlation between the number of pH2A.X foci and the extent of ecDNA untethering (extended Fig 2j-m). This suggests that it is unlikely HDAC inhibition untethers ecDNA via inducing DSBs.

Together, these findings suggest that HDAC inhibition untethers ecDNA from chromosomes by decompacting mitotic chromatin.

### Hypotonic conditions and HDAC inhibition lead to mis-segregation of ecDNA into micronuclei

Next, to determine the fate of untethered ecDNA after cells exit mitosis, I live-imaged COLO320DM cells across mitoses in hypotonic 0.5x media (Fig 3a). ecDNA untethering was observed throughout mitosis. Upon completion of mitosis, some of the untethered ecDNA were incorporated into the primary nuclei of the daughter cells, while some were in the cytoplasm forming micronuclei (Fig 3a). This suggests that like lagging chromosomes and acentric chromatin fragments(Krupina et al., 2021; Warecki & Sullivan, 2020), untethered ecDNA are likely to be mis-segregated into micronuclei.

**Figure 3.**
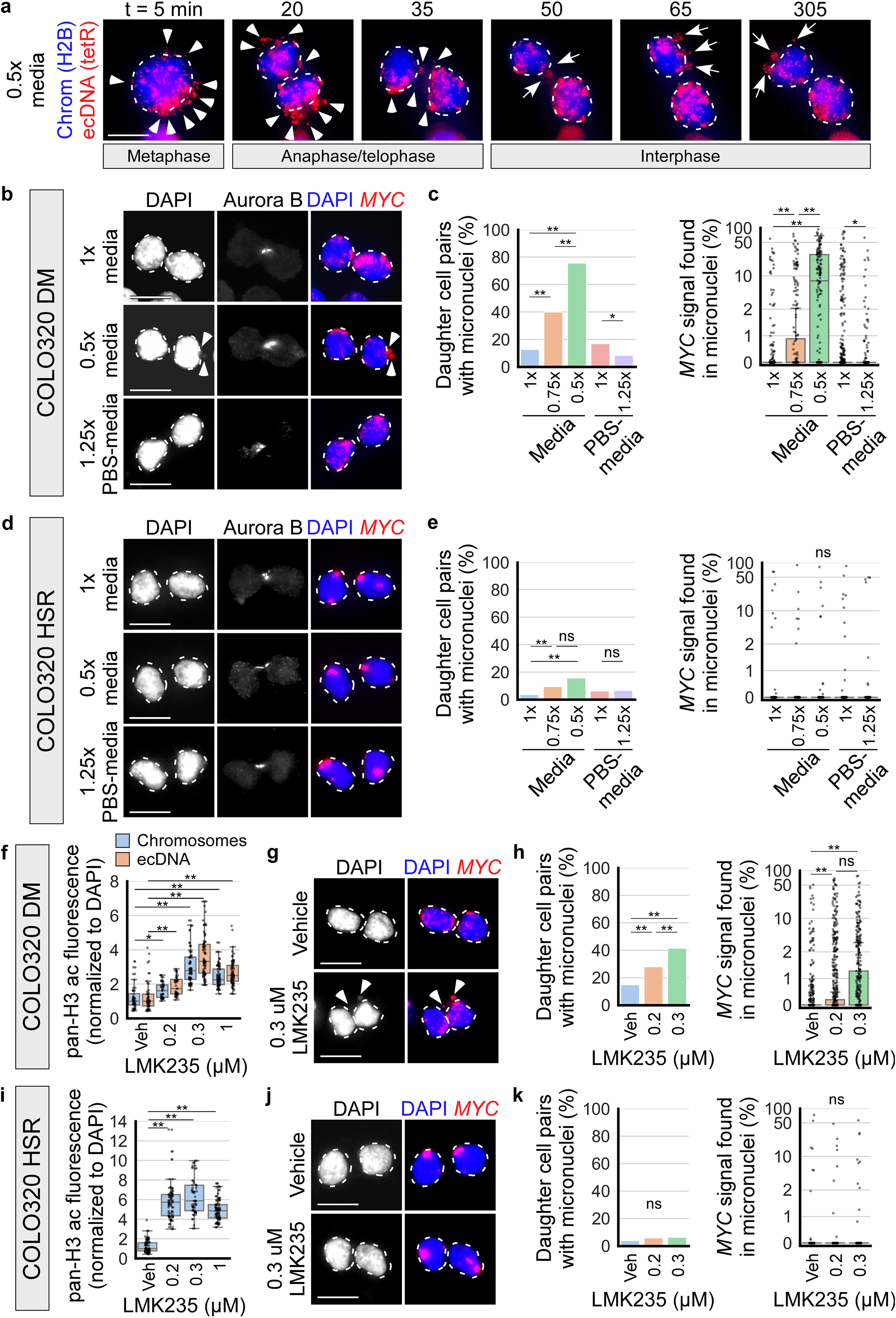
Hypotonic conditions and HDAC inhibition lead to mis-segregation of ecDNA into micronuclei. **a)** Live cell imaging of COLO320DM cells undergoing mitosis (chromatin labeled by H2B-emiRFP670, blue) and (*MYC* loci labeled by tetR-mNeonGreen, red). Mitotic cells were placed in 0.5x media at t=0 min; dashed outlines indicate chromosomes/primary nuclei, arrowheads indicate untethered ecDNA, arrows indicate micronuclei. Representative image of n=5 biological replicates and >50 cells. **b)** Fixed newly-divided daughter COLO320DM cells, as identified by the presence of Aurora B staining, treated for 6 hr with 1x media, 0.75x media (not shown), 0.5x media, 1x PBS-media (not shown), and 1.25x PBS-media; dashed outlines indicate primary nuclei, arrowheads indicate micronuclei. **c)** Quantification of panel b. Left: quantification of the percentage of daughter cell pairs with micronuclei; chi-squared, *p<.05, **p<.01. Right: quantification of the percentage of all *MYC* FISH signal per daughter cell pair inside micronuclei as opposed to the primary nuclei (symlog scale, linear ≤2, log >2); one-way ANOVA, F=151.4, p<.001, *p<.05, **p<.01; n=3, 3, 3, 6, 4 biological replicates and 601, 233, 148, 881, 836 daughter cell pairs. **d/e)** Fixed newly-divided daughter COLO320HSR cells, treated similarly as in panels b/c. **e)** Left: chi-squared, **p<.01, ns = not significant. Right: one-way ANOVA, F=0.9, p=.49; n=3, 3, 3, 5, 3 biological replicates and 680, 256, 179, 578, 487 daughter cell pairs. **f)** Quantification of pan-Histone H3 acetylation (pan-H3ac) IF in cytospin preparations of COLO320DM prometaphase spreads with chromosome and ecDNA segmentation by ecSeg (see extended Fig 3c); cells were treated in vehicle or LMK235 for 24 hr; n=2, 2, 2, 2 biological replicates and 52, 44, 75, 84 cells; one-way ANOVA, chromosomes: F=67.6, p<.001, ecDNA: F=81.3, p<.001, *p<.05, **p<.01 by Tukey’s HSD. **g)** Newly-divided daughter COLO320DM cells, as identified by the presence of Aurora B staining (not shown), treated for 24 hr with vehicle (0.1% DMSO) or indicated concentrations of LMK235; dashed outlines indicate primary nuclei, arrowheads indicate micronuclei. **h)** Quantification of panel g. Left: quantification of the percentage of daughter cell pairs with micronuclei; chi-squared, **p<.01. Right: quantification of the percentage of all *MYC* FISH signal per daughter cell pair inside micronuclei (symlog scale, linear ≤2, log >2); one-way ANOVA, F=151.4, p<.001, *p<.05, **p<.01; n=5, 5, 4 biological replicates and 713, 683, 420 daughter cell pairs. **i)** Same as panel f, for COLO320HSR cells (chromosomes only; see extended Fig 3e); n=2, 2, 2, 2 biological replicates and 85, 85, 63, 99 cells; one-way ANOVA, F=204.1, p<.001, **p<.01 by Tukey’s HSD. **j/k)** Newly-divided daughter COLO320HSR cells, treated similarly as in panels g/h. **k)** Left: chi-squared. Right: one-way ANOVA, F=0.1, p=.88; n=4, 3, 3 biological replicates and 486, 383, 335 daughter cell pairs. In all panels, scale bar = 10 µm.

To quantify the incorporation of untethered ecDNA into micronuclei, I examined micronuclei formation in fixed cells. I treated COLO320DM cells with 1x media, 0.75x media, 0.5x media, 1x PBS-media, and 1.25x PBS-media for 6 hr to allow cells time to undergo mitosis under these conditions. I then fixed and identified newly divided daughter cells based on the presence of Aurora B kinase IF signal between the two daughters, as described previously(Lange et al., 2022) (Fig 3b). I found that the percentage of daughter cell pairs containing micronuclei dramatically increased in cells incubated in hypotonic 0.75x (39.9%) and 0.5x (75.7%) media compared to 1x media (12.6%) (Fig 3b,c). In addition, the percentage of ecDNA (*MYC* FISH signal) within daughter cell pairs found inside micronuclei as opposed to the primary nuclei significantly increased in cells incubated in hypotonic 0.75x (4.2% ± 11.3, mean ± standard deviation) and 0.5x (16.5% ± 21.0) media compared to 1x media (0.6% ± 4.2) (Fig 3c). Furthermore, the micronuclear contents were mainly ecDNA, as measured by the percentage of DAPI signal inside micronuclei that overlapped with *MYC* FISH signal (extended Fig 3a). On the contrary, hypertonic treatment in 1.25x PBS-media decreased ecDNA incorporation into micronuclei compared to 1x PBS-media (Fig 3b,c, extended Fig 3a).

To assess whether hypotonic conditions may induce micronuclei formation in cells that do not contain ecDNA, I compared *MYC* FISH signal incorporation into micronuclei between COLO320DM cells, in which *MYC* is amplified on ecDNA, and COLO320HSR cells, which are derived from the same parental cell line as COLO320DM cells, but in which *MYC* is amplified on chromosomes as homogeneously staining regions (HSR).

The percentage of daughter cell pairs containing micronuclei increased in COLO320HSR cells incubated in hypotonic 0.75x (9.4%) and 0.5x (15.6%) media compared to 1x media (3.5%), although not to the extent seen in COLO320DM cells (Fig 3d,e). However, there was no significant increase in the percentage of *MYC* FISH signal incorporated into micronuclei in COLO320HSR cells incubated in hypotonic 0.75x (0.6% ± 5.8) and 0.5x (1.0% ± 7.2) media compared to 1x media (0.3% ± 3.9) (Fig 3e), and the micronuclei that were formed mostly did not contain *MYC* FISH signal (extended Fig 3b). This indicates that hypotonic conditions may affect the segregation of chromosomes or non-ecDNA acentric fragments. These results suggest that untethered ecDNA are more prone to mis-segregation into micronuclei.

Next, I asked whether HDAC inhibition induces ecDNA micronuclei formation in newly divided daughter cells. TSA treatment was highly cytostatic and prevented cells from completing mitosis; however, at low concentrations, LMK235 and Belinostat treatment allowed cells to progress through mitosis. I treated COLO320DM cells with low concentrations of LMK235 (≤0.3 µM) and Belinostat (≤0.5 µM), which increased histone H3 acetylation in both mitotic chromosomes and ecDNA (Fig 3f and extended Fig 3c). Both inhibitors increased micronuclei formation (41% of daughter cell pairs treated with 0.3 µM LMK235 contained micronuclei compared to 14.7% with vehicle; 58.7% with 0.5 µM Belinostat treatment compared to 11.8% with vehicle) and the percentage of ecDNA incorporated into micronuclei (2.6% with 0.3 µM LMK235 treatment compared to 0.6% with vehicle; 3.0% with 0.5 µM Belinostat treatment compared to 0.7% with vehicle) (Fig 3g,h and extended Fig 3g,h). The micronuclei formed mainly contained ecDNA (extended Fig 3d,i). On the contrary, COLO320HSR cells treated with the same concentrations of LMK235 showed histone hyperacetylation (Fig 3i and extended Fig 3e) but did not exhibit increased micronuclei formation (Fig 3j,k, extended Fig 3f). These results suggest that ecDNA untethering by histone hyperacetylation promotes ecDNA mis-segregation into micronuclei, similarly to the effects observed under hypotonic conditions. This further supports a link between chromatin compaction and ecDNA tethering and segregation.

### Ki67 regulates ecDNA tethering to chromosomes

Ki67 is a large nuclear protein that localizes to the surface of chromosomes during mitosis, forming a surfactant-like molecular layer that prevents chromosomes from tightly clustering together(Cuylen et al., 2016).

I hypothesized that Ki67 may modulate ecDNA tethering similarly to its role in chromosome clustering. First, I assessed the association of Ki67 with mitotic ecDNA by colocalizing Ki67 IF signal with *MYC* FISH signal in COLO320DM cells at metaphase (Fig 4a). To facilitate colocalization analysis, cells were incubated for 15 min in hypotonic 0.5x media to untether ecDNA. *MYC* FISH signal co-localized with Ki67 IF and a line profile across ecDNA shows that Ki67 signal surrounds *MYC* FISH signal (Fig 4b), similar to what was shown for Ki67 and chromosomes in prior studies(Booth et al., 2014; Cuylen et al., 2016; Saiwaki et al., 2005).

**Figure 4.**
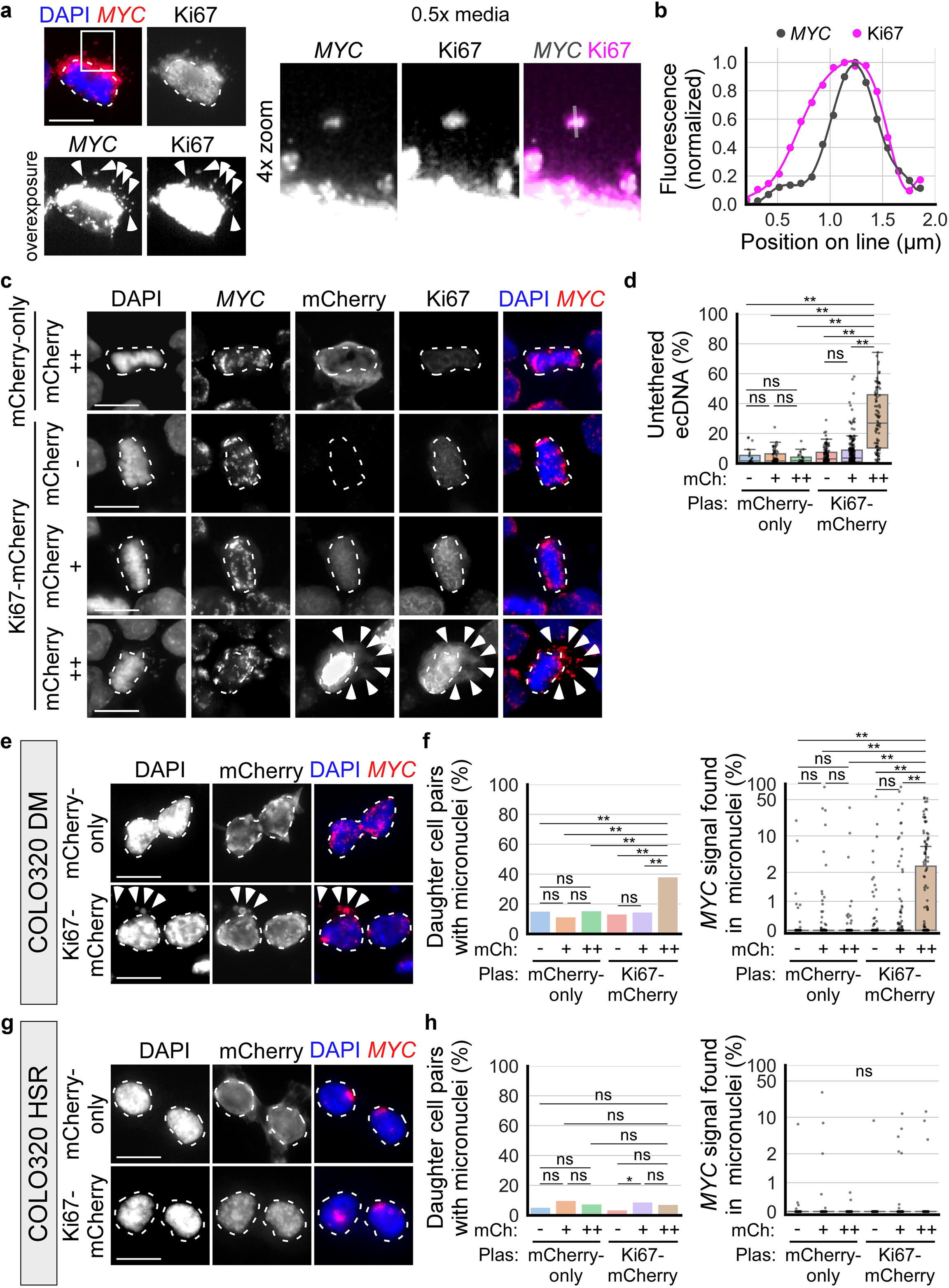
Ki67 gain-of-function untethers ecDNA from chromosomes. **a)** Fixed COLO320DM cells cultured on coverslip and incubated in 0.5x media for 15 min to visualize and colocalize individual ecDNA (*MYC* FISH) with Ki67 IF (arrowheads). **b)** Line profile of *MYC* FISH and Ki67 IF intensity along the line indicated in panel a; polynomial curves were fitted to the line profiles. **c)** Metaphase COLO320DM cells cultured on glass coverslips two to four days post mCherry-only or Ki67-mCherry expression plasmid transfection; cells were categorized based on mCherry fluorescence: - indicates lack of mCherry expression, ++ indicates top 20%tile mCherry expression by fluorescence intensity, + indicates the remaining cells that express mCherry; dashed outlines indicate chromosomes aligned at the metaphase plate, arrowheads indicate untethered ecDNA. **d)** Boxplots quantifying ecDNA untethering in plasmid transfected cells from panel c; mCh = mCherry expression category, Plas = plasmid transfected; from left to right; n=4, 4, 4, 4, 4, 4 biological replicates and 20, 63, 22, 116, 221, 86 total cells; two-way ANOVA, mCherry expression category: F=123.9, p<.001; plasmid transfected: F = 40.5, p<.001; interaction: F=33.8, p<.001; **p<.01 by Tukey’s HSD, ns = not significant. **e)** Newly-divided daughter COLO320DM cells, as identified by the presence of Aurora B staining (not shown), two to four days post mCherry-only or Ki67-mCherry expression plasmid transfection (mCherry ++); dashed outlines indicate primary nuclei, arrowheads indicate micronuclei. **f)** Quantification of panel e. Left: quantification of the percentage of daughter cell pairs with micronuclei; chi-squared, **p<.01. Right: quantification of the percentage of all *MYC* FISH signal per daughter cell pair inside micronuclei (symlog scale, linear ≤2, log >2); two-way ANOVA, mCherry expression category: F=7.6, p<.001; plasmid transfected: F=7.6, p=.0058; interaction: F=4.9, p=.0073; **p<.01; n=7, 7, 7, 7, 7, 7 biological replicates and 93, 312, 104, 179, 400, 147 daughter cell pairs. **g/h)** Newly-divided daughter COLO320HSR cells, transfected similarly as in panels e/f (mCherry ++). **h)** Left: chi-squared, *p<.05. Right: two-way ANOVA, mCherry expression category: F=0.7, p=.49; plasmid transfected: F=0.1, p=.77; interaction: F=0.9, p=.39; n=4, 4, 4, 4, 4, 4 biological replicates and 168, 245, 107, 202, 227, 111 daughter cell pairs. In all panels, scale bar = 10 µm.

Overexpression of Ki67 has been shown to increase the height of the brush-like structure formed by Ki67 at the surface of mitotic chromosomes, further dispersing chromosomes from each other(Cuylen et al., 2016). To test whether Ki67 overexpression may untether ecDNA, I transfected plasmid constructs expressing mCherry-tagged Ki67 (Ki67-mCherry) into COLO320DM cells. At two to four days post transfection, I imaged metaphase cells cultured and fixed on glass coverslips and categorized them based on the level of mCherry expression: mCherry-, indicating no detectable expression; mCherry ++, representing the top 20% in fluorescence intensity; and mCherry +, indicating the remaining mCherry-expressing cells. ecDNA untethering at metaphase was significantly increased in mCherry ++ cells overexpressing Ki67-mCherry (29.0% ± 20.3 untethered, mean ± standard deviation) compared to mCherry ++ cells overexpressing mCherry without Ki67 (mCherry-only; 3.0% ± 3.7) and all other conditions (Fig 4c,d). The percentage of daughter cell pairs containing micronuclei was also increased in cells expressing high levels of Ki67-mCherry (44.9%), compared to mCherry-only (15.4%) (Fig 4e,f). The percentage of ecDNA incorporated into micronuclei was similarly increased (4.6% ± 10.8 in the Ki67-mCherry condition compared to 0.5% ± 3.6 in the mCherry-only condition) in newly divided daughter cells (Fig 4f). The micronuclei formed in cells expressing high levels of Ki67-mCherry mainly contained ecDNA (extended Fig 4a). On the contrary, no increase in micronuclei formation was observed in COLO320HSR cells expressing high levels of Ki67-mCherry (Fig 4g,h, extended Fig 4b). These results show that Ki67 overexpression untethers ecDNA from chromosomes and promotes their mis-segregation into micronuclei.

As a control for protein overexpression, which may affect the intracellular tonicity, I transfected COLO320DM cells with a plasmid construct expressing mCherry-tagged human albumin (mCherry-Albumin), from which the signal peptide sequence is removed, keeping the protein intracellular. Metaphase cells expressing high levels of mCherry-Albumin did not show increased ecDNA untethering (extended Fig 4c,d) nor increased micronuclei formation (extended Fig 4e).

Prior studies have shown that the surfactant function of Ki67 and its ability to disperse mitotic chromosomes do not reside within a single domain of the protein, but rather are due to the size and charge of the protein which, as it coats the surfaces of chromosomes, produces a combination of electrostatic repulsion and steric hindrance preventing inter-chromosomal interactions(Cuylen et al., 2016; Saiwaki et al., 2005). Ki67 is amphiphilic. The Ki67 C-terminal LR (leucine-arginine rich) domain is highly positively charged and preferentially associates with the surface of mitotic chromatin. The remainder of the protein, including the N-terminal forkhead-associated (FHA) domain, the protein phosphatase 1–binding domain (PP1-BD), and the central region contains 16 repeat domains, associates with the cytosol and prevents tight clustering of chromosomes due to its overall positive charge and large, extended structure(Cuylen et al., 2016; Saiwaki et al., 2005). To determine if the ability of Ki67 overexpression to untether ecDNA is similarly dependent on the size of the non-LR domain regions of Ki67 rather than a specific domain, I ectopically expressed in COLO320DM cells one of five truncated versions of Ki67 tagged with mNeonGreen (mNG) as previously described(Cuylen et al., 2016), to determine whether they are able to untether ecDNA. These truncated versions are: i) Ki67-ΔFHA, which lacks the FHA domain, ii) Ki67-ΔNterm, which lacks the FHA and PP1-BD domains, iii) Ki67-ΔRepeats, which lacks the 16 repeat domains, iv) Ki67-LR+8Repeats, which lacks the FHA, PP1-BD and 8 repeat domains, and v) Ki67-LR-only, which lacks all domains except the LR domain (extended Fig 4f). As a control for protein overexpression, I also expressed the Ki67-LR-only construct fused to intracellular albumin (Albumin-Ki67-LR). Similar to prior findings with chromosome dispersal(Cuylen et al., 2016), the expression of Ki67-ΔFHA, Ki67-ΔNterm, and Ki67-ΔRepeats mutants led to ecDNA untethering in COLO320DM metaphase cells and promoted micronuclei formation (extended Fig 4g-h). The expression of smaller Ki67 truncated mutants (Ki67-LR+8Repeats and Ki67-LR-only) and the Albumin-Ki67-LR construct did not increase ecDNA untethering or micronuclei formation (extended Fig 4g-h). These results suggest that Ki67 promotes chromosomal dispersal and ecDNA untethering in similar ways that are dependent on its size and charge more than on specific functional domains.

I next investigated the consequences of Ki67 loss-of-function. I deleted *MKI67* (encoding Ki67) in COLO320DM cells using CRISPR-Cas9 gene editing with one of two guide RNAs (gRNA #1 and gRNA #2) and isolated clones (extended Fig 5a,b). Both gRNAs effectively decreased Ki67 protein levels as assessed by western blotting and IF, compared to control cell clones transfected with a non-targeting guide (NT) (extended Fig 5c,d). Loss of Ki67 slightly but not statistically significantly increased ecDNA tethering (decreased ecDNA untethering) in metaphase COLO320DM cells under isotonic (1x media) conditions (6.0% ± 1.1 untethered with gRNA #1 and 6.7% ± 2.1 with gRNA #2, compared to 7.9% ± 1.5 with NT), indicating that loss of Ki67 is not sufficient to increase ecDNA tethering, at least based on findings on imaging (Fig 5a,b). Loss of Ki67 significantly decreased eDNA untethering in metaphase cells incubated for 15 min in hypotonic 0.75x media (13.7% ± 4.9 untethered with gRNA #1 and 16.5% ± 4.1 with gRNA #2, compared to 28.4% ± 5.0 with NT). Interestingly, the effect was not observed in cells incubated in hypotonic 0.5x media (40.3% ± 8.1 untethered with gRNA #1 and 40.1% ± 5.2 with gRNA #2, compared to 39.6% ± 3.0 with NT), suggesting that stronger hypotonic conditions can compensate for the loss of Ki67, promoting a similar amount of ecDNA untethering in Ki67-deleted cells as in control cells (Fig 5b). Similar results were observed in COLO320DM cells in which *MKI67* expression was knocked down with one of two small interfering RNAs (siRNAs; siMKI67 #1 and #2) compared to a non-targeting control (siControl) (extended Fig 5e-g). In addition, treatment of Ki67-deleted clones with 0.2 µM of the HDAC inhibitor LMK235 for 24 hr did not lead to as much ecDNA untethering as wildtype clones (10.2% ± 0.5 untethered with gRNA #1 and 7.5% ± 0.4 with gRNA #2, compared to 13.3% ± 0.6 with NT) (Fig 5c,d). As with hypotonic conditions, treatment with a higher dose of LMK235 (0.5 µM) compensated for the loss of Ki67 (32.7% ± 7.8 untethered with gRNA #1 and 38.2% ± 6.7 with gRNA #2, compared to 38.3% ± 5.4 with NT), promoting ecDNA untethering equally in control and Ki67-deleted cells (Fig 5d). Altogether, these results indicate that the chromosome surfactant Ki67 coats ecDNA and regulates ecDNA tethering to chromosome (Fig 5e).

**Figure 5.**
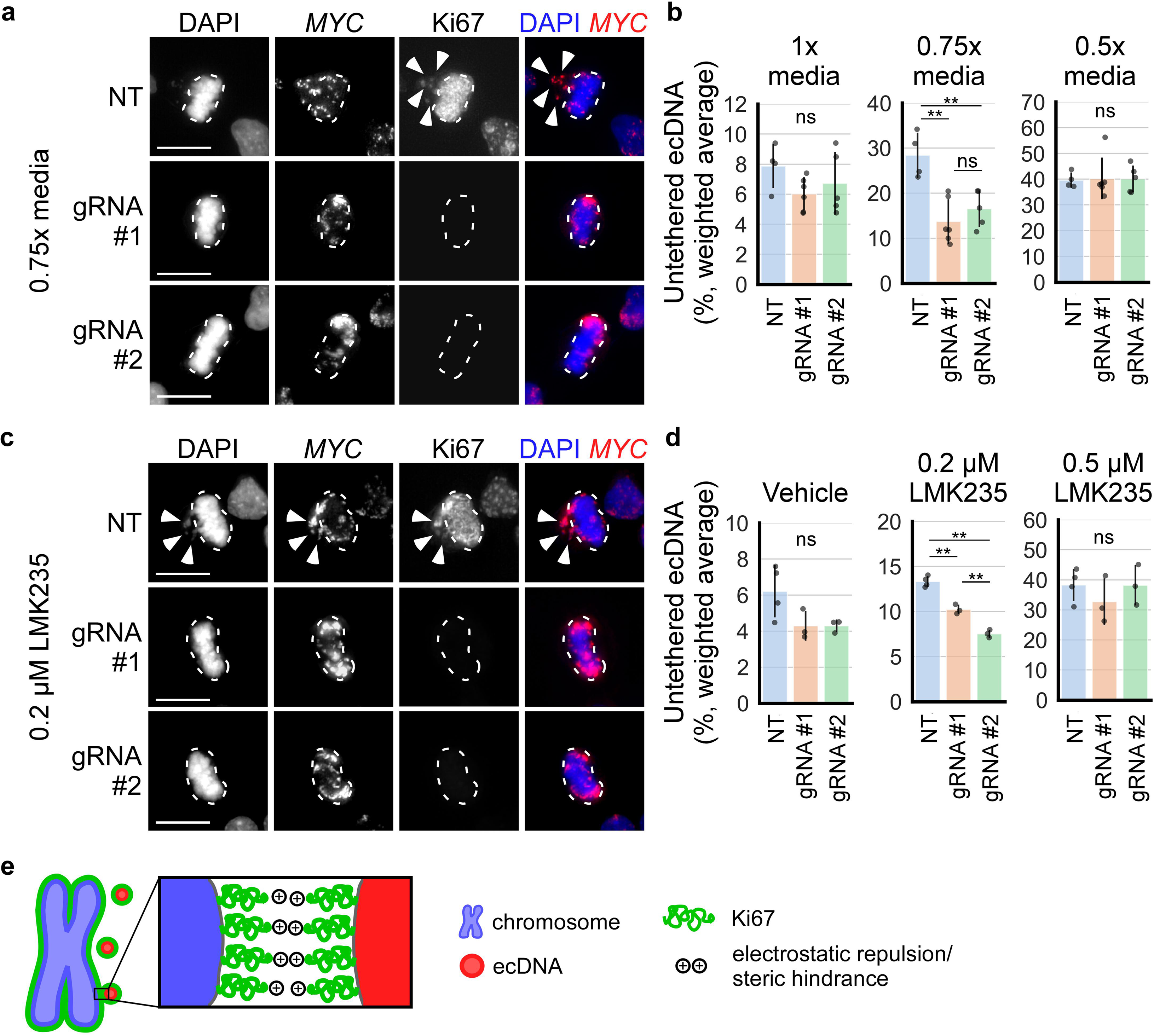
Ki67 loss-of-function decreases ecDNA untethering from chromosomes. **a)** Metaphase COLO320DM wildtype (NT) and *MKI67* knockout cells (gRNA #1 and #2), incubated for 15 min in 1x media (not shown), 0.75x media, and 0.5x media (not shown); dashed outlines indicate chromosomes aligned at the metaphase plate, arrowheads indicate untethered ecDNA. **b)** Quantification of ecDNA untethering of COLO320DM wildtype and *MKI67* knockout clones from panel a (each dot represents the weighted average of at least 95 cells from each clone) after 15 min incubation in 1x media (left; n=4, 6, 5 clones; one-way ANOVA, F=1.7, p=.23), 0.75x media (middle; n=4, 6, 5 clones; one-way ANOVA, F=12.4, p=.001), and 0.5x media (right; n=4, 6, 5 clones; one-way ANOVA, F=0.1, p=.99; **p<.01 by Tukey’s HSD, ns = not significant; error bars = mean ± standard deviation. **c)** Metaphase COLO320DM wildtype (NT) and *MKI67* knockout cells (gRNA #1 and #2), treated for 24 hr with vehicle (not shown), 0.2 µM LMK235, and 0.5 µM LMK235 (not shown); dashed outlines indicate chromosomes aligned at the metaphase plate, arrowheads indicate untethered ecDNA. **d)** Quantification of ecDNA untethering of COLO320DM wildtype and *MKI67* knockout clones from panel c (each dot represents the weighted average of at least 60 cells from each clone) after 24 hr treatment with vehicle (0.1% DMSO, left; n=4, 3, 3 clones; one-way ANOVA, F=3.9, p=.073), 0.2 µM LMK235 (middle; n=4, 3, 3 clones; one-way ANOVA, F=100.8, p<.001, **p<.01 by Tukey’s HSD), and 0.5 µM LMK235 (right; n=4, 3, 3 clones; one-way ANOVA, F=0.8, p=.51); error bars = mean ± standard deviation; *MKI67* knockout clones are gRNA #1 clones 1, 4, and 5, and gRNA #2 clones 1, 4, and 5 (see extended Fig 5a-c). **e)** Schematic: the biological surfactant Ki67 (green) coats the surface of mitotic ecDNA (red) and chromosomes (blue), helping to prevent tethering by electrostatic repulsion and steric hindrance. In all panels, scale bar = 10 µm.

### HDAC inhibition leads to a loss of oncogene copy number in COLO320DM cells, but not in COLO320HSR cells

Next, I asked whether ecDNA untethering and incorporation into micronuclei may affect the inheritance of ecDNA over the course of several cell divisions, leading to a decrease in ecDNA copy number over time. I treated COLO320DM cells, which have a doubling time of roughly 24 hr(Bazzocco et al., 2015), with 1 uM of the HDAC inhibitor LMK235 and quantified ecDNA copy number via *MYC* DNA qPCR after 1 and 2 days of treatment. I observed a 27% reduction in *MYC* copy number at day 2, before cells stopped dividing and/or died. A similar reduction in *MYC* copy number (10-20%) was observed at lower concentrations of LMK235 (0.2 and 0.3 µM) up to 30 days of treatment (Fig 6a). No loss of *MYC* copy number was observed in COLO320HSR cells treated with LMK235 (Fig 6b). Rather, there appeared to be a 10-20% increase in *MYC* copy number at day 10 of treatment. Overall, these data suggest that HDAC inhibition decreases ecDNA copy number.

**Figure 6.**
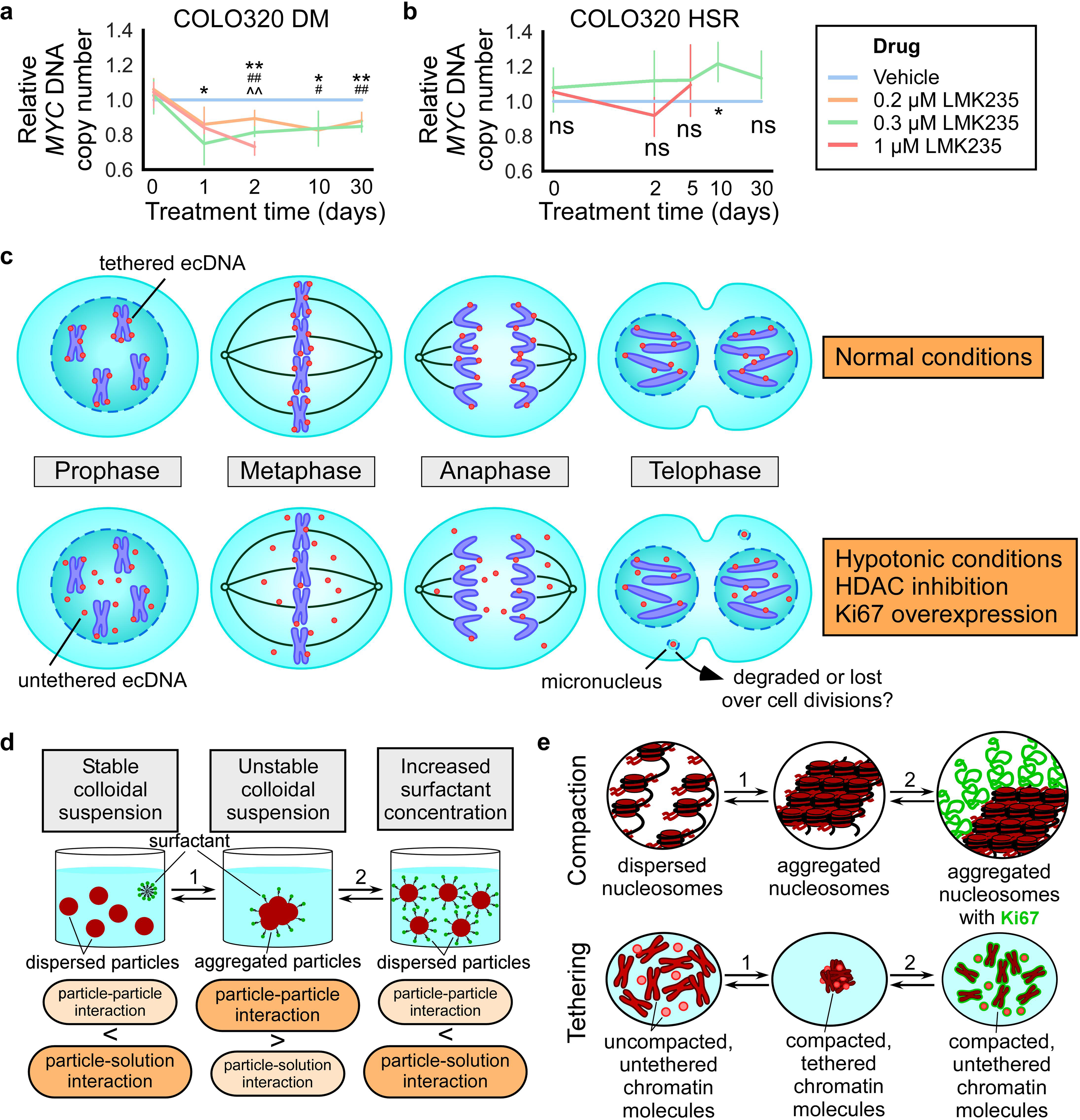
**HDAC inhibition leads to a loss of oncogene copy number in COLO320DM cells, but not in COLO320HSR cells a/b**) Quantification of *MYC* DNA expression in COLO320DM (a) and COLO320HSR (b) cells treated with LMK235; qPCR, 2^-ΔΔCT^ analysis (normalized to *LINE1* copy number and vehicle control); x-axis = symlog scale, linear ≤2, log >2. Error bars = mean ± 95% confidence interval. Drug-containing media was replaced every 2-3 days. **a)** Treatment with vehicle (0.1% DMSO), 0.2 µM, 0.3 µM, or 1 µM LMK235 for 1 day (one-way ANOVA, F=4.1, p=.036, n=4, 4, 4, 3), 2 days (F=31.3, p<.001, n=4, 4, 4, 3), 10 days (F=5.4, p=.017, n=6, 6, 6, no 1 µM), 30 days (F=19.8, p=.0023, n=3, 3, 3, no 1 µM); 0 day: pretreatment (one-way ANOVA, F=0.4, p=.77, n=3, 3, 3, 3); #p<.05, ##p<.01 (0.2 µM LMK235 vs. vehicle); *p<.05, **p<.01 (0.3 µM LMK235 vs. vehicle); ^^p<.01 (1 µM LMK235 vs. vehicle) by Tukey’s HSD; **b)** Treatment with vehicle, 0.3 µM LMK235, or 1 µM LMK235 for 2 days (one-way ANOVA, F=2.2, p=.17, n=4, 4, 4), 5 days (F=0.5, p=.62, n=4, 4, 4), 10 days (F=10.1, p=.019, n=4, 4), and 30 days (F=2.7, p=.15, n=4, 4); 0 day: pretreatment (one-way ANOVA, F=0.8, p=.48); *p<.05 (0.3 µM LMK235 vs. vehicle). **c)** Schematic, top: under normal mitotic conditions, ecDNA tether to chromosomes throughout mitosis to ensure their segregation into primary nuclei of the divided daughter cells; bottom: under conditions that decompact chromatin (hypotonic conditions and HDAC inhibition) or prevent ecDNA-chromosome interaction at their surfaces (Ki67 overexpression), ecDNA untether from chromosomes, leading some to be mis-segregated into micronuclei. **d,e)** Schematics of a colloidal and surface chemistry-based framework for approaching ecDNA and chromosome compaction and tethering during mitosis. Arrow 1 represents biophysical changes to the colloidal particles (nucleosomes and chromatin molecules) or the solution/suspension medium (cytosol), such as alterations to the intracellular ionic strength or to the acetylation state of chromatin. Arrow 2 represents changes in the surfactant (Ki67) concentration of the system. Both sets of changes affect particle-particle and particle-solution interactions.

## Discussion

The mechanisms that tether ecDNA to chromosomes during mitosis and safeguard them from mis-segregation into micronuclei is not completely known(Ilić et al., 2022). Mitotic chromosomes are formed in part by global chromatin compaction driven by increased free intracellular Mg^2+^ concentration and histone deacetylation that modulate chromatin-chromatin interactions(Spicer & Gerlich, 2023). Consequently, mitotic chromatin can be decondensed by hypotonic conditions and HDAC inhibition(Cimini et al., 2003; Kruhlak et al., 2001; Maeshima et al., 2018; Schneider et al., 2022; Wilkins et al., 2014). Here, I show that ecDNA-chromosome tethering is impaired by hypotonic conditions and histone hyperacetylation (Fig 6c), suggesting that tethering is coupled to global chromatin compaction during mitotic chromosome formation. I propose that ecDNA and chromosomes tether to each other via chromatin fiber interactions at the surface of the molecules, mediated by electrostatic and/or hydrophobic forces that also drive mitotic chromatin compaction.

The surfactant Ki67 coats the surface of mitotic chromosomes to help disperse them during mitosis(Booth et al., 2014; Cuylen et al., 2016; Saiwaki et al., 2005; Takagi et al., 2018). I show that Ki67 also associates with ecDNA during mitosis and overexpression of Ki67 untethers ecDNA from chromosomes. Furthermore, Ki67 depletion negates the ecDNA untethering effects of conditions known to decompact mitotic chromatin, i.e. incubation in hypotonic 0.75x media and HDAC inhibition via low dose (0.2 µM) LMK235 (Fig 5a-d). Interestingly, I found an “epistatic” interaction between hypotonic conditions, histone hyperacetylation by HDAC inhibition, and Ki67, as a strong hypotonic condition (0.5x media) and high concentrations of LMK235 (0.5 µM) can compensate for the lack of Ki67, producing high levels of ecDNA untethering despite the lack of Ki67. This “epistatic” interaction show that global chromatin compaction and Ki67-mediated chromatin surface regulation, two of the mechanisms known to help form and organize mitotic chromatin(Spicer & Gerlich, 2023), act together to coordinate ecDNA tethering. I propose that Ki67 disrupts ecDNA-chromosome tethering by preventing chromatin-chromatin interactions at the surface of the molecules during mitotic chromatin compaction.

The finding that ecDNA tethering is coupled to mitotic chromatin compaction suggests that tethering-specific protein complexes and mechanisms are not necessarily required for the proper mitotic segregation of acentric chromatin. I propose that the same tethering mechanisms play a role in the segregation of various forms acentric chromatin, including acentric chromosomal fragments and chromatinized viral DNA, which will be topics of future studies. I hypothesize that the size of the acentric molecules may be a factor in determining whether tethering to chromosomes ensures proper nuclear segregation, as larger molecules experience more drag during mitotic movements in the cytosol, which may act in an opposing direction to tethering forces. This may explain why pieces of chromatin much larger than a single ecDNA, such as lagging chromosomes and large aggregates of ecDNA formed due to DSB, often untether and mis-segregate into micronuclei under normal mitotic conditions(Cimini, 2023; Shimizu et al., 2007; Tanaka & Shimizu, 2000).

DNA inside micronuclei have been suggested to be degraded or lost over several cell divisions (Snapka & Varshavsky, 1983; Terradas et al., 2010; Von Hoff et al., 1992). I found that HDAC inhibition, which untethers ecDNA leading to their mis-segregation into micronuclei, also reduces ecDNA copy number over time (Fig 6a,b). This is an interesting sequelae of HDAC inhibition and while it suggests that ecDNA within micronuclei may be lost over cell divisions due to replication/repair defects or degradation(Di Bona & Bakhoum, 2024), more studies are required to determine the mechanism of the copy number loss.

Both HDAC inhibition(Eckschlager et al., 2017) and modulation of the extracellular tumor environment via hypotonic solutions(Shiozaki et al., 2017) have been used or are potential methods for cancer treatment. I show here that while these treatments may help rid cells of ecDNA, they also cause ecDNA mis-segregation into micronuclei. As mis-segregation and micronuclei formation have been shown to be significant contributors to cancer genomic heterogeneity, progression, and metastasis(Di Bona & Bakhoum, 2024), the costs and benefits of treatments involving HDAC inhibition and hypotonic solutions on ecDNA-containing cancers should be carefully considered and further studied.

Finally, I propose a framework for approaching mitotic chromosome and ecDNA tethering based on colloidal and surface chemistry (Fig 6d). In a stable colloidal suspension of solid particles in a liquid medium, such as a liquid solution, particles are dispersed within the solution, as the particle-particle interaction is weaker than the particle-solution interaction(Hiemenz, 1997). However, upon changes to the biophysical properties of the particles or the solution (arrow 1), the colloidal suspension may become unstable, where particle-particle interactions are stronger than particle-solution interactions. This causes the particles to aggregate, maximizing particle-particle interactions while minimizing particle-solution interactions. Amphiphilic surfactants that stabilize the particle-solution interface help prevent particle aggregation. An increase in surfactant concentration (arrow 2) can disperse the aggregated particles(Hiemenz, 1997). Chromatin suspended in the mitotic cytosol can be modeled as a colloidal suspension at two levels: 1) the level of nucleosomes and 2) the level of chromatin molecules, i.e. chromosomes and ecDNA (Fig 6e). Nucleosomes of decompacted chromatin are dispersed in the cytosol (Fig 6e, top). However, upon an increase in the cytosolic salt/ion concentration or histone deacetylation by HDACs (arrow 1), nucleosomes aggregate, leading to chromatin compaction. This process can be reversed by hypotonic conditions and HDAC inhibition. Zooming out (Fig 6e, bottom), decompacted mitotic chromosomes and ecDNA are dispersed in cytosol, but upon changes that lead to chromatin compaction (arrow 1), they tether to each other due to the same biophysical forces responsible for nucleosome aggregation occurring at the surface of the molecules. This process can be similarly reversed to untether the chromatin molecules. The surfactant Ki67 is large (∼90 nm in length) and acts at the surface of chromatin molecules to stabilize the chromatin-cytosol interface (Cuylen et al., 2016). An increase in the concentration of Ki67 (arrow 2) can untether the chromatin molecules. This framework argues that a complex biological phenomenon such as ecDNA tethering and mitotic inheritance can at least be partially explained by a relatively simple biophysical model of the molecules and their surrounding milieu.

## Materials and Methods

### Cell lines and cell culture

Human colorectal cancer cell lines COLO320DM and COLO320HSR were purchased from ATCC. COLO320DM and HSR cell lines were derived from colorectal carcinoma(Alitalo et al., 1983). Cells were cultured in Dulbecco’s Modified Eagle’s Media with 4.5 g/L Glucose, L-glutamine, and Sodium Pyruvate (Corning) supplemented with 10% fetal bovine serum (Gibco A5256701). All the cells were maintained at 37 °C in a humidified incubator with 5% CO_2_. Cell lines routinely tested negative for mycoplasma contamination. For experiments examining ecDNA untethering at metaphase, cells were treated for 3 hr with 10 µM of the proteasome inhibitor MG132 to enrich for the proportion of cells in metaphase(Wee et al., 2022).

### Chemicals and inhibitors

MG132 (Selleck S2619)

Colcemid (Gibco 15210-040)

Trichostatin A (MedChemExpress HY-15144)

Belinostat (MedChemExpress HY-10225)

Abexinostat (PCI-24781, MedChemExpress HY-10990)

Fimepinostat (CUDC-907, MedChemExpress HY-13522)

Pracinostat (SB939, MedChemExpress HY-13322)

LMK235 (MedChemExpress HY-18998)

TMP269 (MedChemExpress HY-18360)

Mocetinostat (MGCD0103, MedChemExpress HY-12164)

Entinostat (MS275, Tocris 6208)

RGFP966 (Cayman 16917)

Santacruzamate A (CAY10683, Cayman 15403)

Tasquinimod (MedChemExpress HY-10528)

BRD73954 (MedChemExpress HY-18700)

Nicotinamide (Cayman 11127)

EX527 (Cayman 10009798)

CTPB (Cayman 19570)

5-Azacytidine (Sigma Aldrich A2385)

GSK343 (Selleck S7164)

RRx001 (Selleck S8405)

ZM447439 (Selleck S1103)

Hesperadin (Selleck S1529)

Paprotrain (Selleck E0779)

DAPI (Invitrogen D1306)

### Antibodies

Rabbit anti-H3ac, pan-acetyl (ActiveMotif 39140); used at 1:200 for IF Rabbit anti-phospho-H2A.X (Cell Signaling S139); used at 1:250 for IF Rabbit anti-Aurora B kinase (Bethyl A300-431A); used at 1:200 for IF Mouse anti-Aurora B kinase (Novus NBP2-50039); used at 1:2000 for IF Mouse anti-Ki67 (eBioscience 14-5699-82); used at 1:500 for IF, 1:1000 for WB Rabbit anti-mCherry (abcam 167453); used at 1:500 for IF Mouse anti-mNeonGreen (Proteintech 32f6); used at 1:500 for IF Mouse anti-Actin (Sigma A4700); used at 1:10000 for WB Anti-mouse IgG and anti-rabbit secondary antibodies, Alexa Fluor (Invitrogen); used at 1:500 for IF Anti-mouse IgG, HRP-linked (Cell Signaling 7076S); used at 1:10000 for WB

### Plasmids

Plasmids were transfected into cells using Lipofectamine 3000 Transfection Reagent (Thermo L3000001) following manufacturer’s instructions.

Ki67-ΔFHA expression: Ki67ΔFHA-mNeonGreen-IRESpuro2 was a gift from Daniel Gerlich (Addgene plasmid # 183739; http://n2t.net/addgene:183739; RRID:Addgene_183739). Ki67-ΔNterm expression: Ki67ΔNterm-mNeonGreen-IRESpuro2 was a gift from Daniel Gerlich (Addgene plasmid # 183740; http://n2t.net/addgene:183740; RRID:Addgene_183740). Ki67-ΔRepeats expression: Ki67Δrepeats-mNeonGreen-IRESpuro2 was a gift from Daniel Gerlich (Addgene plasmid # 183742; http://n2t.net/addgene:183742; RRID:Addgene_183742). Ki67-LR+8Repeats expression: Ki67 (8 repeats+LR)-mNeonGreen-IRESpuro2 was a gift from Daniel Gerlich (Addgene plasmid # 183741; http://n2t.net/addgene:183741; RRID:Addgene_183741). Ki67-LR-only expression: Ki67 (LR)-mNeonGreen-IRESpuro2 was a gift from Daniel Gerlich (Addgene plasmid # 183743; http://n2t.net/addgene:183743; RRID:Addgene_183743).

New plasmid constructs were generated using Gibson assembly. Briefly, assembly fragments were generated either by restriction digest or PCR amplification with Q5 high-fidelity DNA polymerase (New England Biolabs M0491), following manufacturer’s protocols. Gibson assembly was performed using NEBuilder HiFi DNA Assembly Reaction Master Mix (New England Biolabs E2621) following manufacturer’s protocols. The assembly reactions were used to transform One Shot Stbl3 Chemically Competent Cells (Invitrogen C7373). For Ki67-mCherry expression, Ki67-mCherry-IRESpuro2 plasmid was generated by replacing mNeonGreen with mCherry coding sequence in Ki67-mNeonGreen-IRESpuro2 (a gift from Daniel Gerlich, Addgene plasmid # 183737; http://n2t.net/addgene:183737; RRID:Addgene_183737). For mCherry-only expression, mCherry-IRESpuro2 plasmid was generated by replacing Ki67-mNeonGreen with mCherry coding sequence in Ki67-mNeonGreen-IRESpuro2 pIRESpuro2-mCherry. For mCherry-Albumin expression, mCherry-albumin-IRESpuro2 plasmid was generated by fusing human serum albumin coding sequence (from pGEM-Alb, SinoBiological HG10968-G) without signal peptide downstream of mCherry (removing TAA from mCherry) in mCherry-IRESpuro2. For Albumin-Ki67-LR expression, Albumin-Ki67(LR)-mNeonGreen-IRESpuro2 plasmid was generated by fusing human serum albumin coding sequence without the signal peptide in-frame upstream of Ki67-LR in Ki67(LR)-mNeonGreen-IRESpuro2.

### *MKI67* siRNA knockdown

Predesigned Dicer-Substrate siRNAs from TriFECTa RNAi Kit (IDT hs.Ri.MKI67.13) were used, including non-targeting siRNA (siControl), hs.Ri.MKI67.13.1 (siMKI67 #1), and hs.Ri.MKI67.13.2 dsiRNAs (siMKI67 #2), both targeting exon 13 of *MKI67* (NM_002417). Cells were forward transfected using Lipofectamine RNAiMAX Transfection Reagent (Invitrogen 13778075) at an siRNA concentration of 10 nM, following manufacturer’s instructions. Knockdown efficiency was confirmed by qRT-PCR and IF.

### *MKI67* knockout cell line generation

CRISPR/Cas9 gene editing: predesigned Alt-R CRISPR-Cas9 guide RNAs (gRNA) were used, including Hs.Cas9.MKI67.1.AC (gRNA #1, targeting exon 2 and Hs.Cas9.MKI67.1.AA (gRNA #2), targeting exon 6 *MKI67* (NG_047061.1). Non-targeting gRNA (NT) sequence (GAGAGTGCGCCTTGATAGTA) was selected from a published list(Doench et al., 2016) based on sequence similarity to the targeting gRNAs. Alt-R S.p. Cas9-RFP V3 enzyme (IDT 10008162) and gRNA ribonucleoprotein (RNP) complex was formed following manufacturer’s instructions and forward transfected using Lipofectamine CRISPRMAX (Invitrogen CMAX00003) at a final RNP concentration of 10 nM, following manufacturer’s instructions. Single cell cloning by limiting dilution: at 48 hr post RNP transfection, editing efficiency was confirmed via heteroduplex formation and T7 endonuclease I digestion using the Alt-R Genome Editing Detection Kit (IDT 1075931) following manufacturer’s instructions. PCR primers: *MKI67* exon 2 F 5’-GACTTGACGAGCGGTGGTTC-3’, R 5’-GCACCAAGGAAAAGTGACGG-3’; *MKI67* exon 6 F 5’-TGACCACTAGCTCCCAACTG-3’, R 5’-AGGCATCTGTCTGTCGTTGAC-3’. Cells were diluted to 0.5 cell/100 µL in conditioned media and seeded at 100 µL per well in a 96 well plate. Cells were then monitored for single colony formation. Once wells with single colonies reached 50% confluency, cells were transferred to a 24 well plate. gDNA was harvested from each clone, PCR amplified, and submitted for Sanger sequencing (Azenta Life Sciences) to confirm mutations and clonal purity. Mutations were quantitated using TIDE (https://tide.nki.nl/)(Brinkman et al., 2014). Knockout efficiency was confirmed by IF and Western blotting.

### Quantitative PCR (qPCR and qRT-PCR)

*MYC* copy number enumeration: gDNA was harvested from cells using QIAamp DNA Mini Kit (Qiagen 51304), following manufacturer’s instructions. qPCR was performed using iQ SYBR Green Supermix (Bio-Rad 1708880) following manufacturer’s instructions. *MYC* primers: F 5’-AGCGACTCTGGTAAGCGAAG-3’, R 5’-TGGCCCGTTAAATAAGCTGC-3’. *LINE-1* was used for normalization; primers: F 5’-AAAGCCGCTCAACTACATGG-3’, R 5’-TGCTTTGAATGCGTCCCAGAG-3’. 2^-ΔΔCT^ method was used for analysis.

*MKI67* qRT-PCR: RNA was harvested from cells using RNeasy Mini Kit (Qiagen 74106), following manufacturer’s instructions. cDNA was synthesized using SuperScript VILO cDNA Synthesis Kit (Invitrogen 11754050) following manufacturer’s instructions. qPCR was performed as above. *MKI67* primers: F 5’-CGGTCCCTGAAAATAAGGGAATATC-3’, R 5’-TCTGGGGAGGTCTTCATGGG-3’. *GAPDH* was used for normalization; primers sequences were obtained from a prior publication(Kwon et al., 2021): F 5’-TTGGCTACAGCAACAGGGTG-3’, R 5’-GGGGAGATTCAGTGTGGTGG-3’. 2^-ΔΔCT^ method was used for analysis.

### Western blotting (WB)

Cell lysis: cells were washed with 1x PBS and lysed in 1x RIPA buffer (Thermo 89901) with protease and phosphatase inhibitors (Thermo 1861281) for 30 min on ice with gentle agitation. After centrifugation at 4 C, 14000 x g for 15 min, the supernatant was flash frozen in liquid nitrogen and stored at-80 C until needed. Protein concentration was determined using the Pierce BCA Protein Assay Kit (Thermo 23225). Gel electrophoresis and transfer: cell lysates mixed with Laemmli sample buffer (Bio-Rad 1610747) and 2-mercaptoethanol (Aldrich M6250) and separated on Protean TGX stain free gels (Bio-Rad 456-8123). Trans-blot turbo transfer pack (Biorad 1704159) was used to transfer the protein onto a 0.2 µm nitrocellulose membrane. Antibody staining: after 1 hr blocking in TBS + 0.1% Tween (TBST) with 3% bovine serum albumin, the membrane was incubated in the primary antibody diluted in blocking solution overnight at 4 C with gentle agitation. After washing in TBST, the membrane was then incubated in HRP-conjugated secondary antibody diluted in TBST, washed in TBST, and then visualized using the SuperWest Pico PLUS Chemiluminescent kit (Thermo 34579).

### Prometaphase spreads, drop method

For performing FISH on prometaphase spreads, cells were incubated for 3 hr in 0.1 µg/ml colcemid for 3 hr, then washed with 1x PBS and transferred to Eppendorf tubes. Cells were then resuspended in 75 mM KCl (or the indicated solution) and incubated at 37 C for 15 min with agitation. Cells were then fixed with Carnoy’s fixative (3:1 mix of methanol:glacial acetic acid) added dropwise, followed by three more exchanges of fresh Carnoy’s fixative. Fixed cells were dropped from 10 cm onto a humidified slide and allowed to dry before staining.

### Prometaphase spreads, cytospin method

For immunostaining of prometaphase spreads, cells were incubated for 3 hr in 0.1 µg/ml colcemid (Gibco 15210-040) for 3 hr, then washed with 1x PBS and transferred to Eppendorf tubes. Cells were then gently resuspended in ice-cold hypotonic buffer (10 mM Tris-Cl (pH 7.5) + 20 mM NaCl + 5 mM KCl + 1 mM CaCl_2_ + 0.5 mM MgCl_2_ + 40 mM glycerol) and incubated on ice for 10 min. ∼10k cells were cytospun in 200 µl of hypotonic buffer + 0.25% Tween-20 for each slide at 800 RPM for 8 min.

### Immunofluorescence staining (IF)

Cells cultured on glass coverslips: cells were cultured on glass coverslips coated with 10 µg/mL human plasma fibronectin (Millipore FC010). After fixation in 4% PFA, cells were permeabilized for 10 min in 1x PBS + 0.25% Triton X-100, then blocked for 1 hr in 1x PBS + 3% bovine serum albumin. Cells were then incubated with primary antibody diluted in blocking solution at 4 C overnight. After washing with 1x PBS + 0.1% Tween-20, cells were incubated in fluorescent-dye conjugated secondary antibody diluted in blocking solution. After washing, cells were counterstained with DAPI before mounting.

Cells on cytospin slides: cells fixed in 4% PFA were permeabilized and blocked with DAKO antibody diluent (Agilent S080983-2) + 5 mM MgCl_2_ + 0.2% Triton X-100 for 10 min. Cells were then incubated with primary antibody diluted in blocking solution at 4 C overnight. After washing with blocking solution + 0.1% Tween-20, cells were incubated in fluorescent-dye conjugated secondary antibody diluted in blocking solution. After washing, cells were counterstained with DAPI before mounting.

### Fluorescence in situ hybridization (FISH)

Slides with specimen were dehydrated in an ethanol series (70%, 85%, 100%) and air-dried. Fluorescently-tagged *MYC* probe (Empire Genomics) diluted in hybridization buffer is applied over the specimen and spread using a coverslip. The slides were then warmed to 75 C for 3 min to denature DNA before incubating in a humidified chamber at 37 C overnight. Slides were washed in 0.4x SSC, followed by 2x SSC + 0.1% Tween-20, and then counterstained with DAPI before mounting.

Dual IF-FISH: after IF staining, cells were fixed in 4% PFA and then permeabilized with 1x PBS + 0.7% Triton X-100 + 0.1 M HCl for 10 min. Cells were then incubated in 1.9 M HCl for 30 min to denature DNA. Cells were then dehydrated in an ethanol series (70%, 85%, 100%) and air-dried. FISH probe is diluted in hybridization buffer and heated to 75 C for 3 min, then allowed to cool briefly before being applied over the specimen and spread using a coverslip. Slides are then placed in a humidified chamber at 37 C overnight. Slides were washed in 0.4x SSC, followed by 2x SSC + 0.1% Tween-20, and then counterstained with DAPI before mounting.

### Microscopy

Fluorescence microscopy was performed on a Leica DMi8 widefield microscope by Las X software (v.3.8.2.27713) using a ×63 oil objective. Z stacks were acquired covering the entire thickness of cells, with 0.5 µm step size. For live cell imaging, cells were cultured in FluoroBrite media (Gibco A18967-01) in 96 well glass-bottom plates and imaged on a Leica DMi8 widefield microscope using a ×63 oil objective at 37 C with 5% CO_2_.

### Image analysis

Except where indicated below, all image analysis steps were automated using custom Python scripts with the packages numpy, OpenCV, and scikit-image. Custom graphical user interface (GUI) software for region of interest selection and manual segmentation mask correction were created using the Python package tkinter. All custom scripts are available on github: image processing: https://github.com/yanglum/extract_images_from_lif_file, analyses of prometaphase spread:

https://github.com/yanglum/metaseg_gui, analyses of fixed cells cultured on glass coverslips:

https://github.com/yanglum/ecDNA_stain_analysis_scripts.

<u>Post-acquisition image processing:</u> for prometaphase spread images, the best in-focus z plane was selected for further image analysis based on Laplacian variance. For live cells and fixed cells coated on glass coverslips, image deconvolution using Leica’s Lightning and Thunder processing was performed, and max projection images were made from z-stacks.

<u>ecDNA untethering and chromosome individualization in prometaphase spreads:</u> prometaphase spreads were segmented using the metaseg tool from ecSeg (https://github.com/UCRajkumar/ecSeg)(Rajkumar et al., 2019) to distinguish between ecDNA and chromosomes based on DAPI staining, with manual correction of the segmentation mask, as needed, using MYC FISH to identify tethered ecDNA. Connected component (CC) analysis was then performed to quantify ecDNA and chromosome count per prometaphase cell. The percentage of ecDNA untethering is calculated as the number of untethered ecDNA CCs divided by the total number of ecDNA CCs (untethered ecDNA CCs + tethered ecDNA CCs). Untethered ecDNA is defined as ecDNA CC with <25% of its perimeter adjacent to pixels classified as chromosome. Tethered ecDNA is defined as ecDNA CC with 25% to 99.9% of its perimeter adjacent to chromosome pixels. ecDNA CCs with 100% of its perimeter adjacent to chromosome pixels are not counted. The number of chromosome CCs per prometaphase cell is used to represent the amount of chromosome individualization. For the inhibitor screen in Fig 3c and extended Fig 3a,b, DAPI and *MYC* FISH signal were segmented by Otsu’s thresholding method and the convex hull area was calculated based on the segmentation for each cell. The convex hull area is used as an estimate of ecDNA and chromosome untethering.

<u>pan-H3ac immunofluorescence quantification in prometaphase spreads:</u> prometaphase spreads were segmented using the metaseg tool from ecSeg as above. Average pan-H3ac immune-and DAPI fluorescence intensity were quantified for all pixels classified as chromosome or ecDNA per cell. Pan-H3ac fluorescence is normalized to DAPI fluorescence to control for DNA content (ecDNA have lower DAPI staining intensity than chromosomes).

<u>ecDNA untethering and micronuclei quantification in fixed cells cultured on glass coverslips:</u> cells were segmented for DAPI and MYC FISH signal using Otsu’s thresholding method. For quantification of ecDNA untethering, metaphase cells were manually identified on DAPI based on the appearance of chromosomes aligned at the metaphase plate. The largest DAPI CC was labeled as the genomic mass of chromosomes aligned at the metaphase plate. MYC FISH positive pixels overlapping with the genomic mass were quantitated as tethered, while those not overlapping with the genomic mass were quantitated as untethered. For micronuclei analysis, newly divided daughter cells were manually identified based on the presence of Aurora B kinase IF between the two cells. DAPI segmentation was used as the segmentation mask with the primary nuclei and micronuclei manually identified and labeled. Micronuclei were defined as smaller areas of DAPI signal found outside of the large primary nuclei. For pH2A.X foci quantification, pH2A.X IF signal was segmented using Otsu’s thresholding method and the number of foci per cell was quantified using CC analysis, where CCs consisting of more than five pixels were counted as foci. For IF signal quantification (Ki67, mCherry, mNeonGreen), the IF signal was segmented using Otsu’s thresholding method and the total IF signal (integrated density) per cell was quantified within the segmented region.

### Graphing and statistics

All graphing and statistical analyses were performed in Python using the packages numpy, pandas, matplotlib, seaborn, statsmodels (ANOVA, Tukey’s HSD) and scipy stats (linear regression, Student’s t test, chi-squared). All statistical methods and sample sizes can be found in the figure legends. No statistical methods were used to predetermine the sample size. For ANOVA, Tukey’s honestly significant difference (HSD) was used for posthoc analysis when p<.05.

## Acknowledgements

I thank Dr. Serena Sanulli for her assistance in preparing the manuscript, and I.T.W. and B.R. for their critical reading of the manuscript. I also thank the Sanulli and Mischel labs for their helpful discussions and input. This work was supported by the Department of Pathology at Stanford University School of Medicine and through its Physician Scientist Incubator program.

---

**Extended Figure 1.**
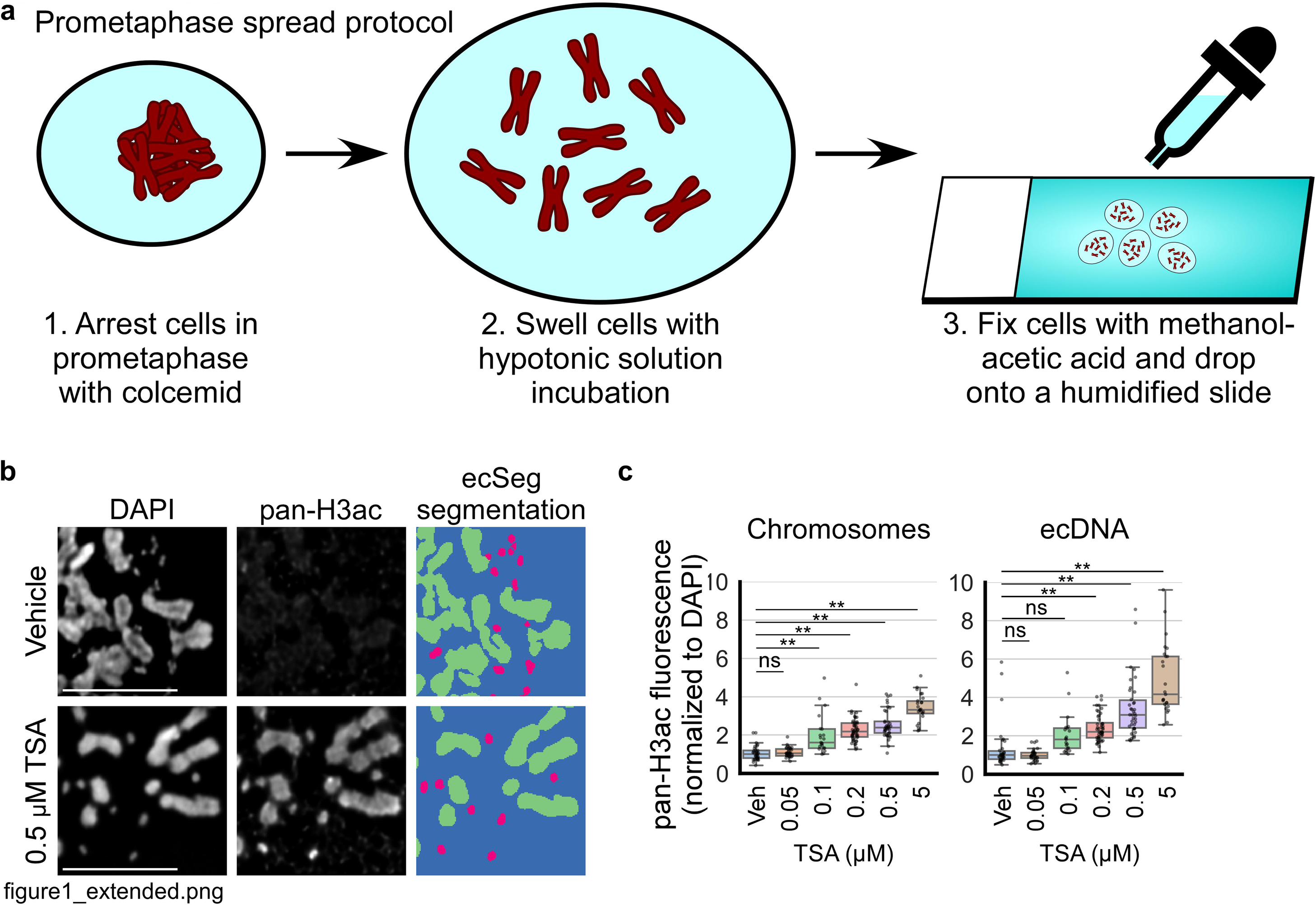
**a)** The prometaphase spread technique involves three main steps. 1) Cells are arrested in prometaphase by treatment with a mitotic spindle destabilizing drug, such as colcemid. 2) Cells are incubated in hypotonic solution, conventionally 75 mM KCl. 3) Cells are fixed with a 3:1 mixture of methanol and acetic acid and dropped onto a humidified slide. **b)** Cytospin preparations of prometaphase-arrested COLO320DM cells incubated in hypotonic solution, with IF staining for pan-Histone H3 acetylation (pan-H3ac). ecSeg was used to classify pixels as either chromosome (green) or ecDNA (red). Scale bar = 10 µm. **c)** Boxplots quantifying pan-H3ac IF signal by integrated density from panel b, normalized to DAPI signal, for chromosomes (left) and ecDNA (right); from left to right, n=3, 1, 1, 1, 2, 2, biological replicates and 34, 33, 21, 48, 38, 25 cells. Left: one-way ANOVA, F=60.8, p<.001. Right: one-way ANOVA, F=44.1, p<.001; **p<.01 by Tukey’s HSD, ns = not significant.

**Extended Figure 2.**
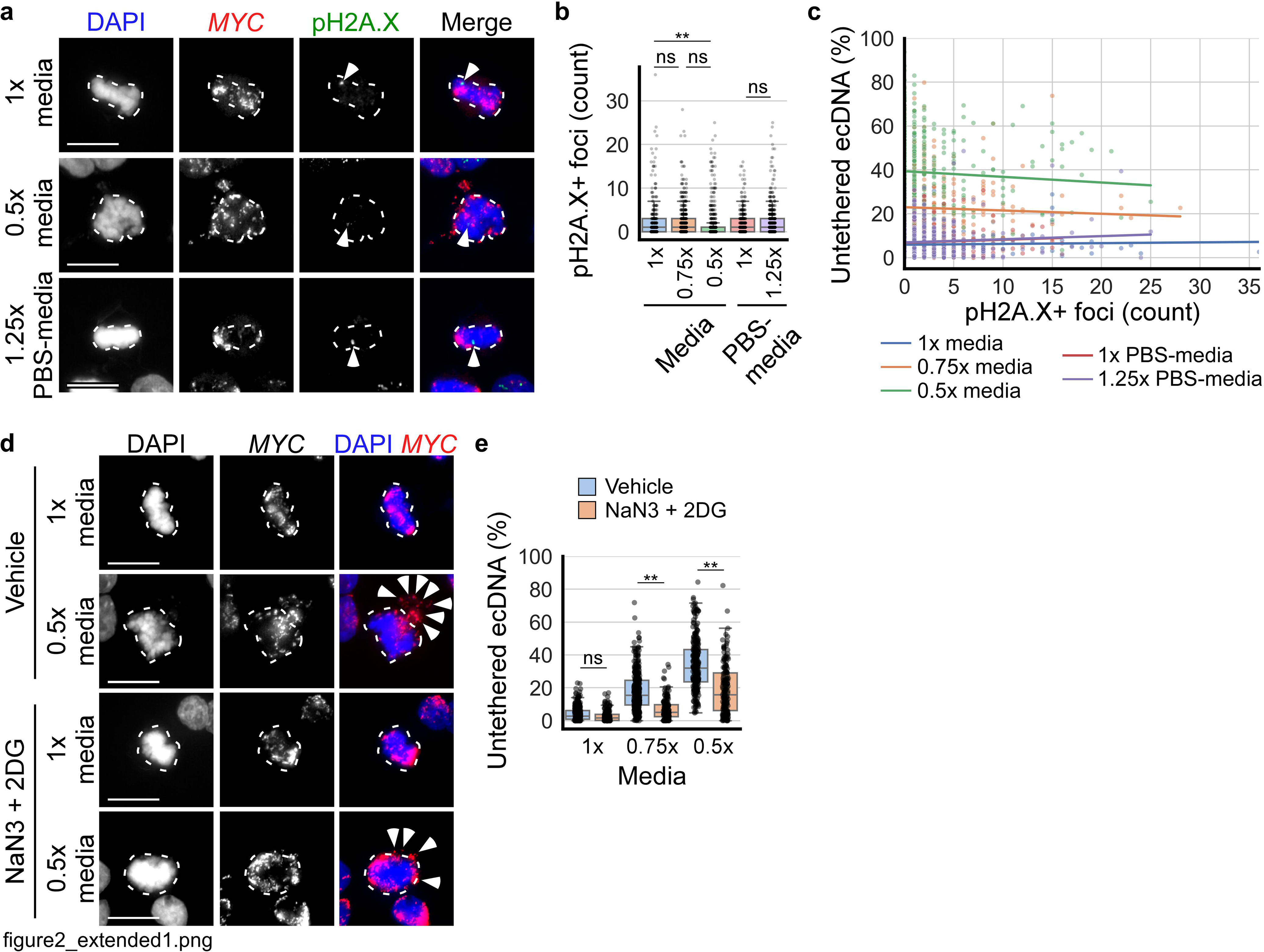
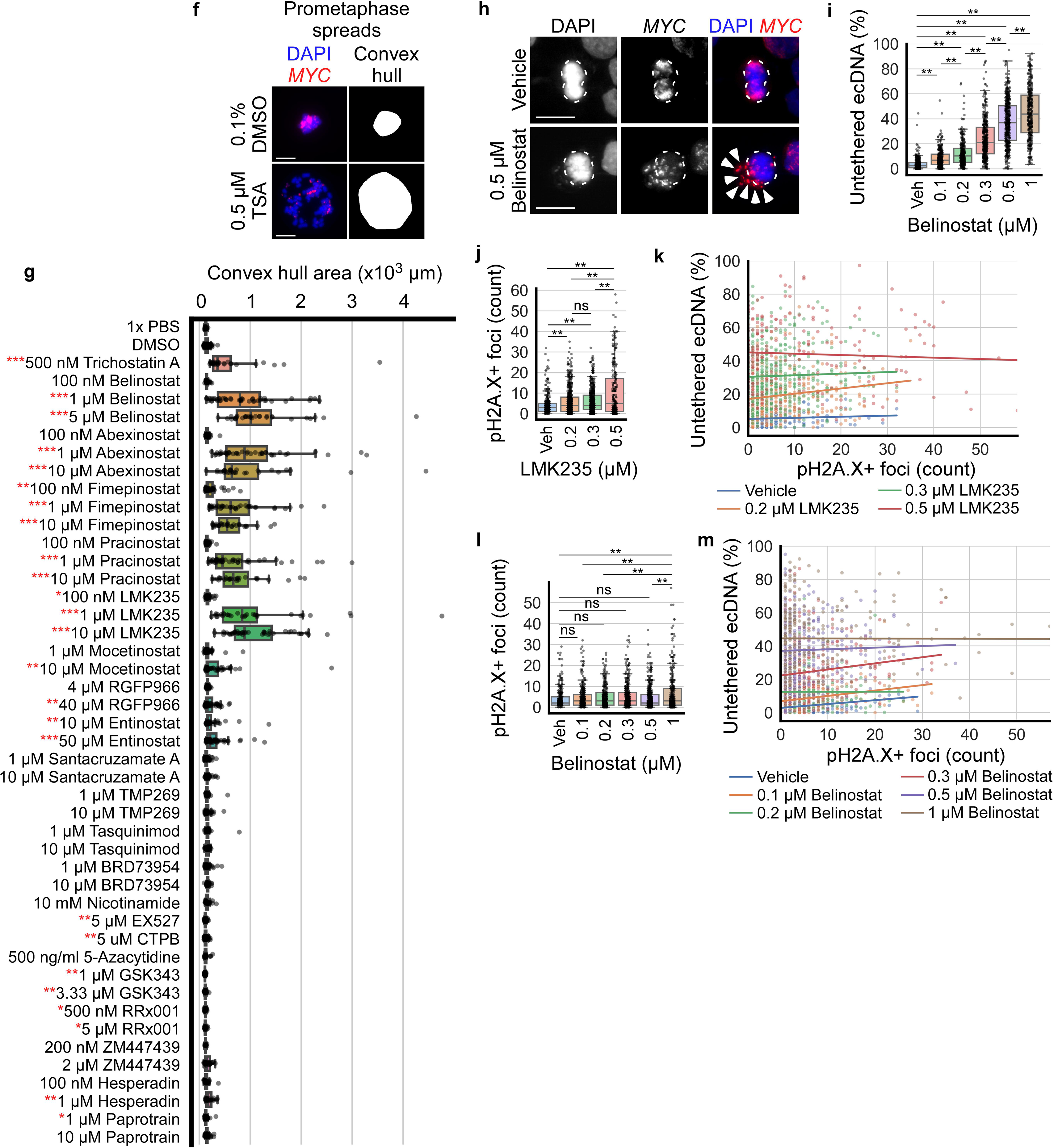
**a)** Metaphase COLO320DM cells treated for 15 min with 1x media, 0.75x media (not shown), 0.5x media, 1:1 mix of 1x PBS with 1x media (1x PBS-media, not shown), and 1:1 mix of 1.5x PBS with 1x media (1.25x PBS-media); dashed outlines indicate metaphase plate chromosomes, arrowheads indicate pH2A.X foci. **b)** Boxplots quantifying the number of pH2A.X foci per metaphase cell from panel a; from left to right, n=3, 3, 3, 3, 3 biological replicates and 304, 515, 576, 338, 506 cells; one-way ANOVA, F=5.4, p<.001, **p<.01 by Tukey’s HSD, ns = not significant. **c)** Correlation of the number of pH2A.X foci with ecDNA untethering for each treatment condition from panel b. **d)** COLO320DM cells arrested in metaphase with 10 µM MG132, treated for 3 hr with vehicle (regular media) or 10 mM NaN_3_ and 50 mM 2-Deoxy-D-glucose (2-DG) dissolved directly in cell culture media and then incubated for 15 min in 1x media, 0.75x media (not shown), or 0.5x media; dashed outlines indicate metaphase plate chromosomes, arrowheads indicate untethered ecDNA. **e)** Quantification of untethered ecDNA per cell from panel d; n=4, 4, 4, 4, 4, 4 biological replicates and 328, 194, 341, 174, 272, 169 cells; two-way ANOVA, NaN3 + 2DG: F=230.9, p<.001; media: F=657.8, p<.001; interaction: F=50.4, p<.001, **p<.01 by Tukey’s HSD, ns = not significant. **f)** Representative images of prometaphase spreads performed without incubation in hypotonic solution, performed on COLO320DM cells treated for 24 hr with 0.1% DMSO and 0.5 µM TSA treatment, along with the convex hull of the spread. **g)** Boxplots quantifying the convex hull area of prometaphase spreads produced without incubation in hypotonic solution after 24 hr treatment with the indicated drug; from top to bottom, n=14, 205, 28, 49, 46, 30, 54, 40, 29, 44, 39, 29, 47, 44, 28, 39, 31, 43, 51, 55, 42, 55, 40, 59, 102, 114, 59, 38, 54, 31, 57, 54, 30, 60, 60, 18, 42, 43, 21, 24, 11, 31, 32, 21, 59, 62; *padj<0.05, **padj<0.001, ***padj<1e-10, padj = adjusted Student’s t-test p value using Bonferroni multiple test correction (44 total comparisons were made). **h)** Metaphase COLO320DM cells cultured on glass coverslips treated for 24 hr with vehicle (0.1% DMSO) or indicated concentrations of Belinostat; dashed outlines indicate metaphase plate chromosomes, arrowheads indicate untethered ecDNA. **i)** Boxplots quantifying untethered ecDNA per cell from panel h; n=3, 3, 3, 3, 3, 3 biological replicates and 291, 310, 276, 357, 430, 316 cells; one-way ANOVA, F=424.3, p<.001, **p<.01 by Tukey’s HSD. **j)** Quantification of the number of pH2A.X foci per metaphase COLO320DM cell treated for 24 hr with vehicle (0.1% DMSO) or indicated concentration of LMK235; from left to right, n=5, 5, 4, 3 biological replicates and 331, 515, 472, 215 cells; one-way ANOVA, F=39.8, p<.001, **p<.01 by Tukey’s HSD. **k)** Correlation of the number of pH2A.X foci with ecDNA untethering for each treatment condition from panel j. **l)** Quantification of the number of pH2A.X foci per metaphase COLO320DM cell treated for 24 hr with vehicle (0.1% DMSO) or indicated concentration of Belinostat; n=3, 3, 3, 3, 3, 3 biological replicates and 291, 310, 276, 357, 430, 316 cells; one-way ANOVA, F=6.2, p<.001, **p<.01 by Tukey’s HSD. **m)** Correlation of the number of pH2A.X foci with ecDNA untethering for each treatment condition from panel l. In all panels, scale bar = 10 µm.

**Extended Figure 3.**
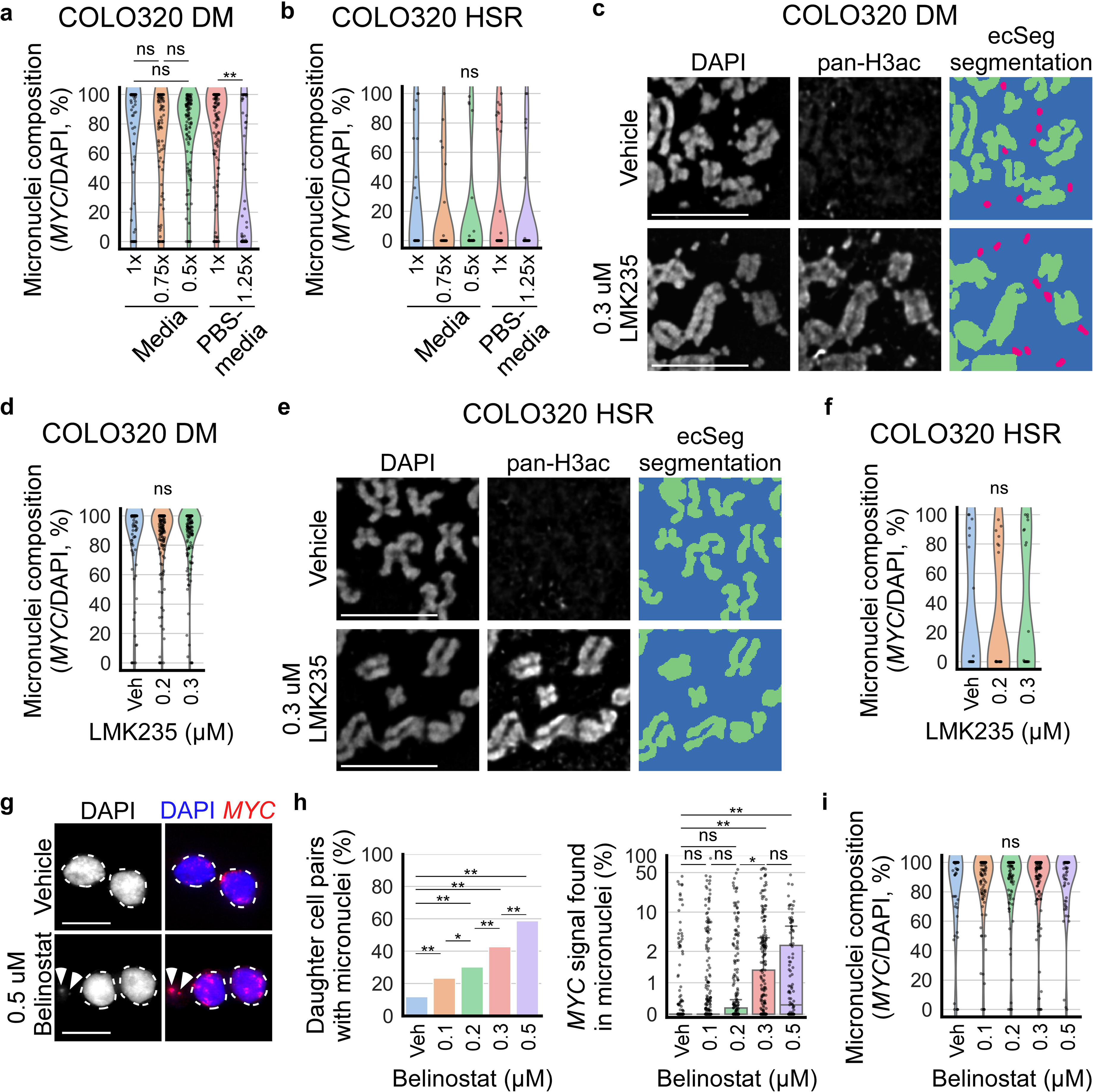
**a)** Violin plots quantifying micronuclei composition (area of *MYC* FISH signal divided by area of DAPI signal within micronuclei) in COLO320DM cells treated for 6 hr with 1x media, 0.75x media, 0.5x media, 1x PBS-media, and 1.25x PBS-media from Fig 3b; n=3, 3, 3, 6, 4 biological replicates and 63, 85, 110, 131, 65 daughter cell pairs (all micronuclei combined per cell pair); one-way ANOVA, F=19.1, p<.001, **p<.01. **b)** same quantification as panel a but for COLO320HSR cells from Fig 3d; n=3, 3, 3, 5, 3 biological replicates and 23, 24, 28, 36, 32 daughter cell pairs; one-way ANOVA, F=0.6, p=.63. **c)** Cytospin preparations of COLO320DM prometaphase spreads treated in vehicle or LMK235 for 24 hr, with IF staining for pan-Histone H3 acetylation (pan-H3ac). ecSeg was used to classify pixels as either chromosome (green) or ecDNA (red). Quantified in Fig 3f. **d)** Violin plots quantifying micronuclei composition in COLO320DM daughter cell pairs treated for 24 hr with vehicle (0.1% DMSO) or indicated concentration of LMK235 from Fig 3g; n=5, 5, 4 biological replicates and 104, 187, 171 daughter cell pairs (all micronuclei combined per cell pair); one-way ANOVA, F=0.1, p=.86. **e)** Same as panel c, for COLO320HSR cells; quantified in Fig 3i. **f)** same quantification as panel d but for COLO320HSR cells from Fig 3j; n=4, 3, 3 biological replicates and 19, 22, 21 daughter cell pairs; one-way ANOVA, F=0.2, p=.80. **g)** Newly-divided daughter COLO320DM cells, as identified by the presence of Aurora B staining, treated for 24 hr with vehicle (0.1% DMSO) or indicated concentrations of Belinostat; dashed outlines indicate primary nuclei, arrowheads indicate micronuclei. **h)** Quantification of panel g. Left: quantification of the percentage of daughter cell pairs with micronuclei; chi-squared, *p<.05, **p<.01. Right: quantification of the percentage of all *MYC* FISH signal per daughter cell pair inside micronuclei (symlog scale, linear ≤2, log >2); one-way ANOVA, F=8.0, p<.001, *p<.05, **p<.01; n=3, 3, 3, 3, 3 biological replicates and 482, 483, 358, 333, 121 daughter cell pairs. **i)** Violin plots quantifying micronuclei composition from panel g; n=3, 3, 3, 3, 3 biological replicates and 54, 107, 104, 135, 68 daughter cell pairs; one-way ANOVA, F=0.6, p=.64. In all panels, scale bar = 10 µm.

**Extended Figure 4.**
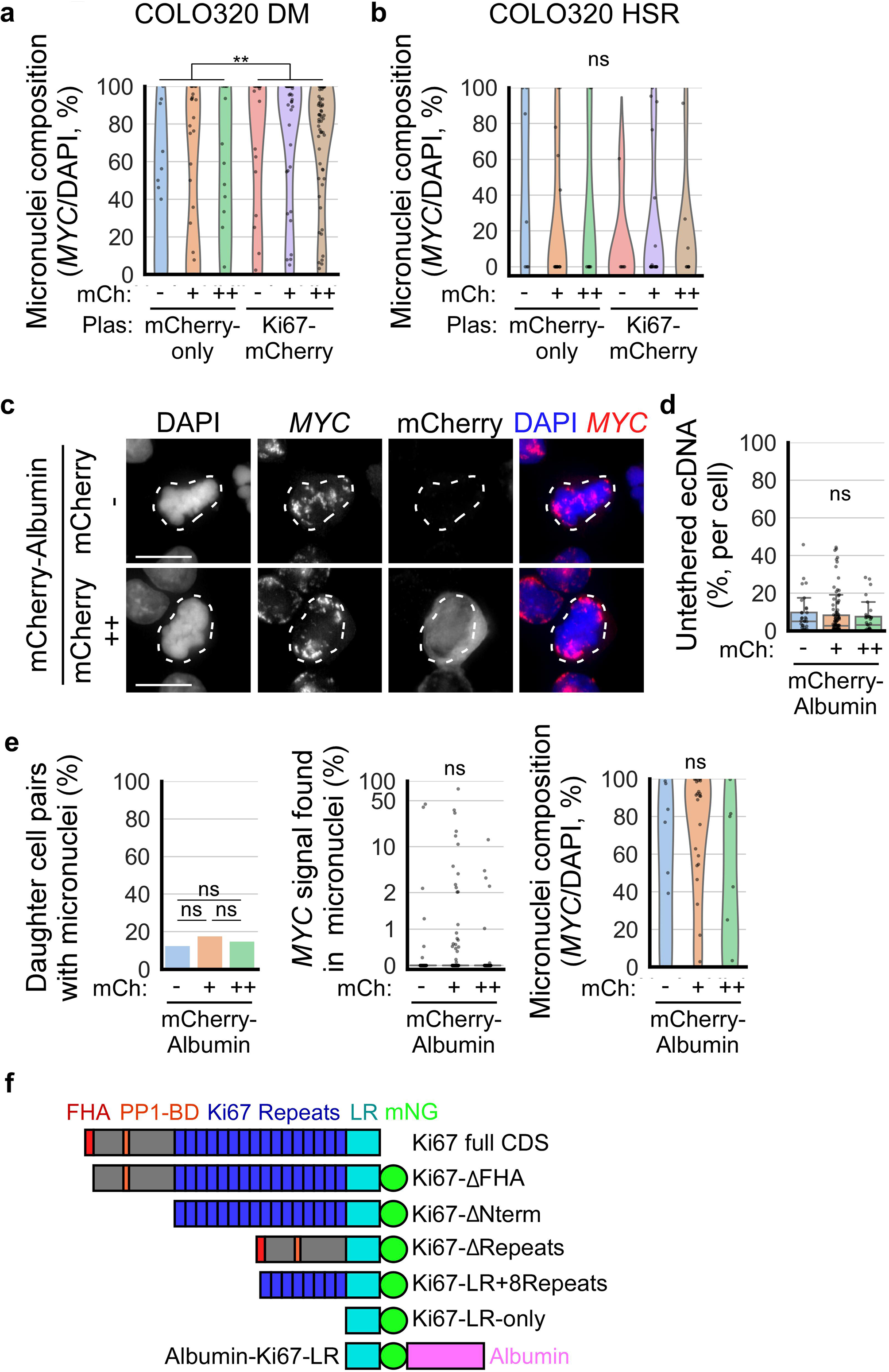
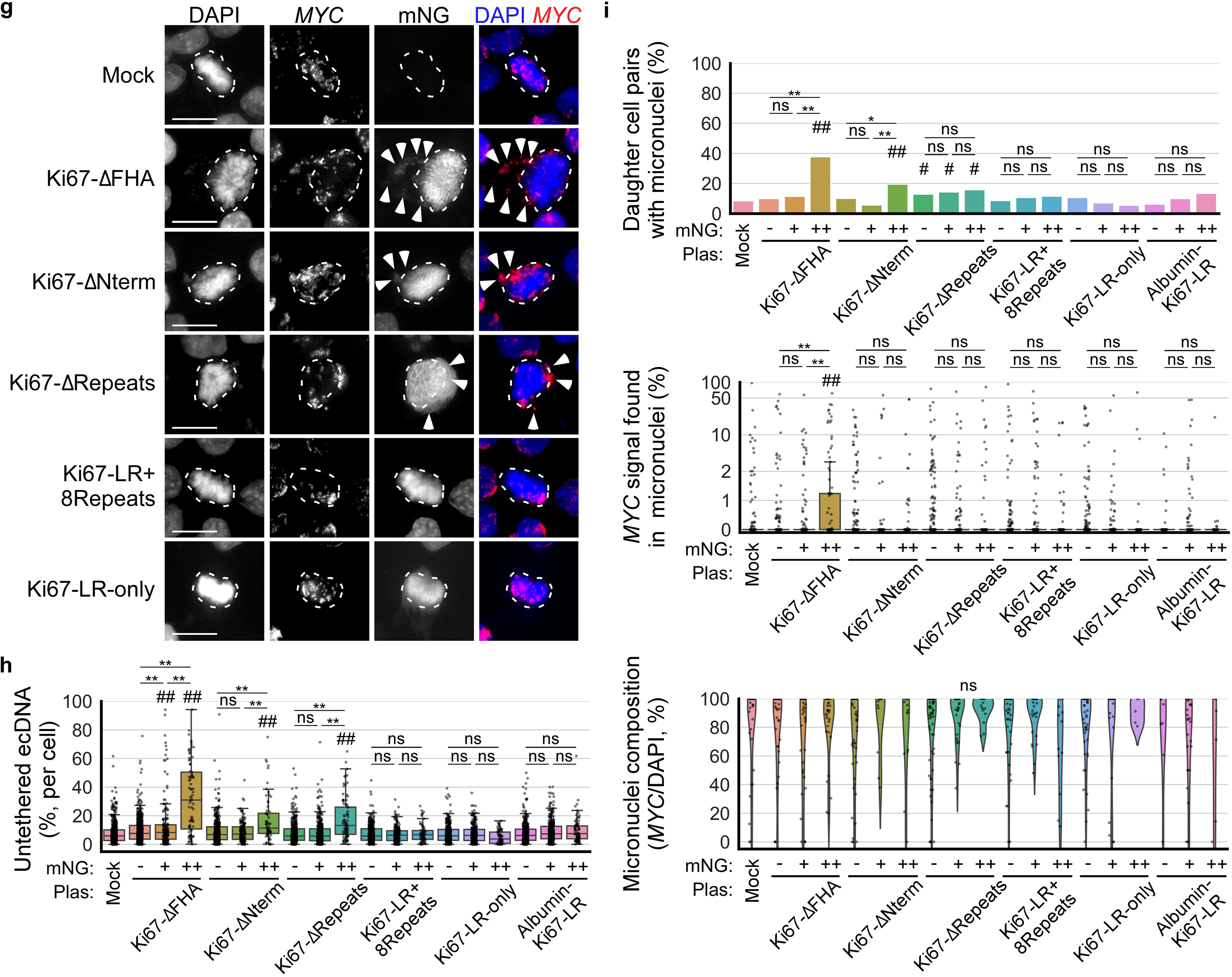
**a)** Violin plots quantifying micronuclei composition in COLO320DM daughter cell pairs two to four days post mCherry-only or Ki67-mCherry expression plasmid transfection from Fig 4e; mCh = mCherry expression category, Plas = plasmid transfected; from left to right, n=7, 7, 7, 7, 7, 7 biological replicates and 13, 31, 15, 23, 53, 55 daughter cell pairs (all micronuclei combined per cell pair); two-way ANOVA, mCherry expression category: F=0.4, p=.70; plasmid transfected: F=9.3, p=.0027; interaction: F=0.0, p=.97, ns = not significant. **b)** same quantification as panel a but for COLO320HSR cells from Fig 4g; n=4, 4, 4, 4, 4, 4 biological replicates and 7, 18, 8, 6, 20, 8 daughter cell pairs; two-way ANOVA, mCherry expression category: F=0.2, p=.83; plasmid transfected: F=0.9, p=.34; interaction: F=0.9, p=.40. **c)** Metaphase COLO320DM cells two to four days post mCherry-Albumin (albumin coding sequence with signal peptide removed) expression plasmid transfection; cells were categorized based on mCherry fluorescence: - indicates lack of mCherry expression, ++ indicates top 20%tile mCherry expression by fluorescence intensity, + indicates the remaining cells that express mCherry; dashed outlines indicate chromosomes aligned at the metaphase plate. **d)** Boxplots quantifying ecDNA untethering in panel c; n=3, 3, 3 biological replicates and 36, 106, 37 cells; one-way ANOVA, F=0.4, p=.70. **e)** Left: quantification of the percentage of COLO320DM daughter cell pairs transfected with mCherry-Albumin plasmid with micronuclei, chi-squared. Middle: quantification of the percentage of all *MYC* FISH signal per daughter cell pair inside micronuclei, one-way ANOVA, F=0.5, p=.64; n=4, 4, 4 biological replicates and 71, 162, 60 daughter cell pairs. Right: quantification of micronuclei composition, one-way ANOVA, F=1.9, p=.17; n=4, 4, 4 biological replicates and 9, 27, 8 daughter cell pairs. **f)** Schematic of Ki67 domains and truncated Ki67 constructs (adapted from(Cuylen et al., 2016)), mNG = mNeonGreen. **g)** Metaphase COLO320DM cells two to four days post transfection with truncated Ki67 constructs; dashed outlines indicate chromosomes aligned at the metaphase plate; arrowheads indicate untethered ecDNA. **h)** Boxplots quantifying ecDNA untethering in panel g; cells were categorized based on mNeonGreen fluorescence: - indicates lack of mNeonGreen expression, ++ indicates top 10%tile mNeonGreen expression by fluorescence intensity, + indicates the remaining cells that express mNeonGreen; n=5, [5, 5, 5], [5, 5, 5], [5, 5, 5], [5, 5, 5], [5, 5, 5], [5, 5, 5] biological replicates and 555, [532, 168, 81], [414, 152, 66], [368, 386, 86], [400, 211, 71], [336, 268, 70], [385, 358, 85] cells; two-way ANOVA, mNeonGreen expression category: F=120.8, p<.001, plasmid transfected: F=39.3, p<.001, interaction: F=34.4, p<.001, **p<.01 for indicated pair-wise comparisons, ##p<.01 compared to mock by Tukey’s HSD. **i)** Quantification of COLO320DM cells two to four days post transfection with truncated Ki67 constructs. Top: quantification of the percentage of COLO320DM daughter cell pairs with micronuclei, chi-squared, *p<.05, **p<.01, #p<.05 compared to mock, ##p<.01 compared to mock. Middle: quantification of the percentage of all *MYC* FISH signal per daughter cell pair inside micronuclei (symlog scale, linear ≤2, log >2), two-way ANOVA, mNeonGreen expression category: F=15.2, p<.001, plasmid transfected: F=1.7, p=.12, interaction: F=3.5, p<.001; n=4, [4, 4, 4], [4, 4, 4], [4, 4, 4], [4, 4, 4], [4, 4, 4], [4, 4, 4] biological replicates and 502, [469, 351, 93], [572, 250, 93], [648, 259, 102], [673, 323, 113], [554, 412, 109], [193, 334, 60] daughter cell pairs. Bottom: quantification of micronuclei composition, two-way ANOVA, mNeonGreen expression category: F=0.5, p=.50, plasmid transfected: F=1.4, p=.21, interaction: F=1.4, p=.18; n=4, [4, 4, 4], [4, 4, 4], [4, 4, 4], [4, 4, 4], [4, 4, 4], [4, 4, 4] biological replicates and 35, [39, 35, 33], [51, 12, 15], [65, 33, 14], [43, 31, 12], [51, 24, 5], [8, 30, 6] daughter cell pairs. In all panels, scale bar = 10 µm.

**Extended Figure 5.**
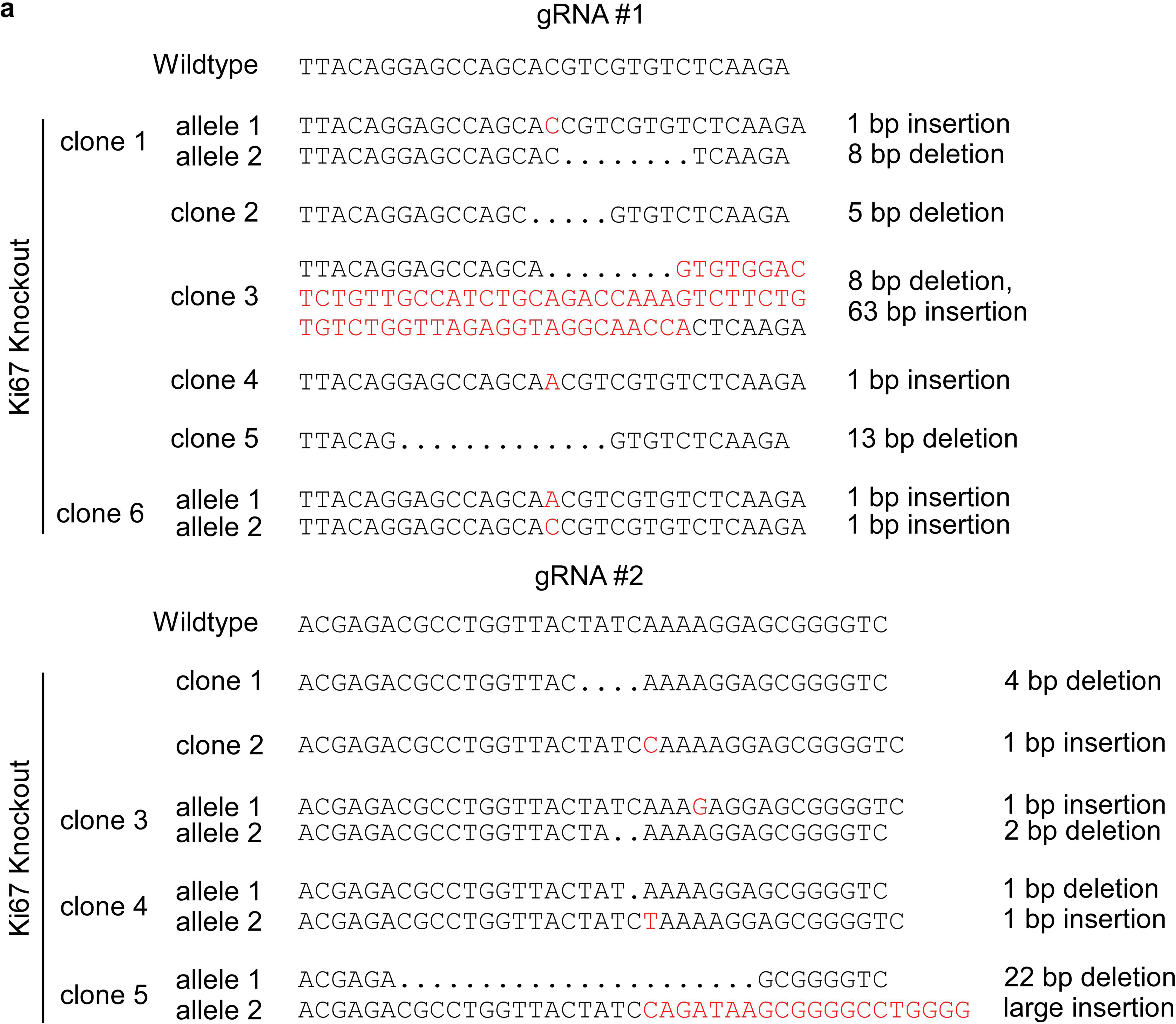
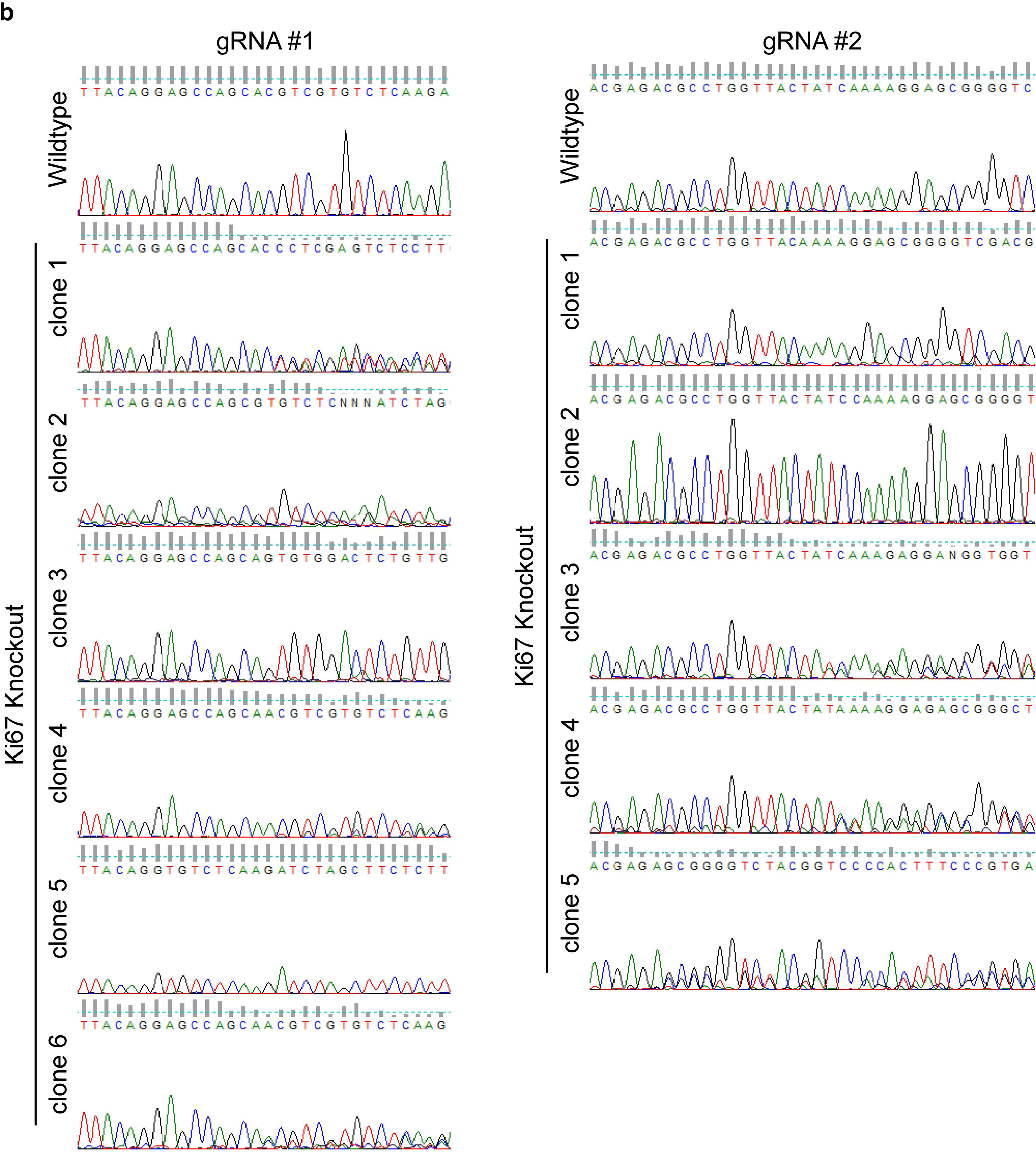
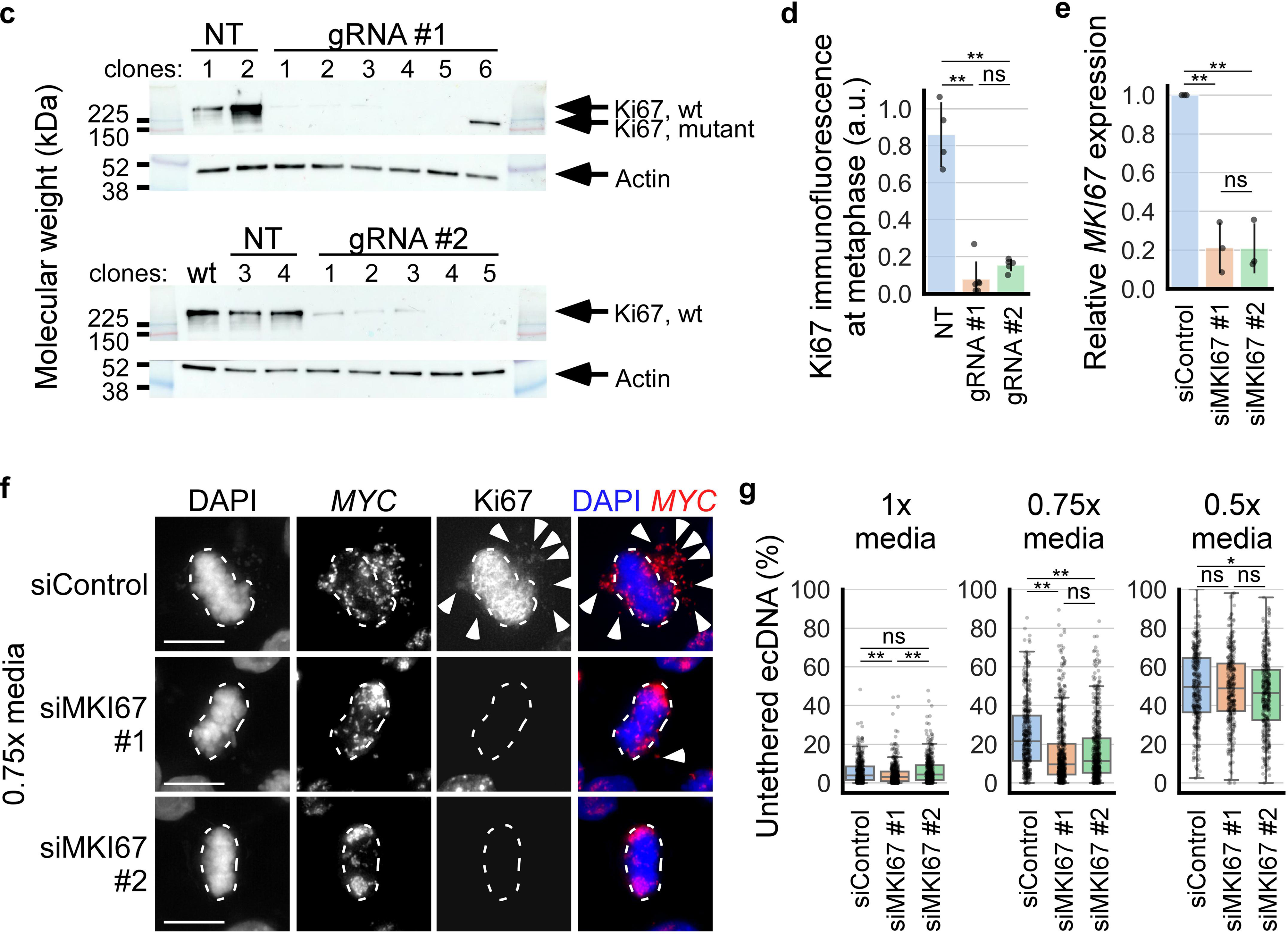
**a)** Indel mutations generated in each *MKI67* knockout clone by CRISPR-Cas9 mediated gene editing of *MKI67* at exon 6 (gRNA #1) and exon 2 (gRNA #2). **b)** DNA sequence chromatograms at the CRISPR-Cas9 target site for each clone. **c)** Western blot performed on whole cell lysates of wildtype COLO320DM cells (wt, not transfected with Cas9) and *MKI67* wildtype and knockout clones generated using CRISPR-Cas9 mediated gene editing with non-targeting (NT) and *MKI67*-targeting gRNAs (gRNA #1 and #2). **d)** Quantification of Ki67 IF signal by integrated density in metaphase COLO320DM wildtype (NT) and *MKI67* knockout clones (gRNA #1 and #2); each dot represents the average of at least 122 cells from each clone; from left to right, n=4, 6, 5 clones; one-way ANOVA, F=70.1, p<.001, **p<.01 by Tukey’s HSD, ns = not significant; error bars = mean ± standard deviation; a.u. = arbitrary units. **e)** Quantification of *MKI67* mRNA expression in COLO320DM cells two days post transfection with non-targeting siRNA (siCtrl) or one of two siRNAs targeting *MKI67* (siMKI67 #1 and #2); qRT-PCR, 2^-ΔΔCT^ analysis (normalized to *GAPDH* and siCtrl); n=3, 3, 3 biological replicates; one-way ANOVA, F=56.2, p<.001, **p<.01 by Tukey’s HSD, ns = not significant; error bars = mean ± standard deviation. **f)** Metaphase COLO320DM cells cultured on glass coverslips two days post siRNA transfection, incubated for 15 min in 1x media (not shown), 0.75x media, and 0.5x media (not shown); dashed outlines indicate chromosomes aligned at the metaphase plate, arrowheads indicate untethered ecDNA; scale bar = 10 µm. **g)** Boxplots quantifying ecDNA untethering in siRNA transfected metaphase COLO320DM cells from panel f after 15 min incubation in 1x media (left; n=3, 3, 3 biological replicates and 419, 454, 477 cells; one-way ANOVA, F=56.2, p<.001), 0.75x media (middle; n=3, 3, 3 biological replicates and 407, 572, 543 cells; one-way ANOVA, F=41.0, p<.001), and 0.5x media (right; n=3, 3, 3 biological replicates and 336, 313, 366 cells; one-way ANOVA, F=3.9, p=.02); *p<.05, **p<.01 by Tukey’s HSD.

